# Optogenetic tripwires resolve models of how cells count DNA breaks

**DOI:** 10.1101/2025.10.31.685943

**Authors:** Marco Labagnara, Vojislav Gligorovski, Sahand Jamal Rahi

## Abstract

DNA end resection is a critical step in DNA damage repair and activation of the DNA damage checkpoint (DDC). To date, resection has been studied primarily in bulk cell populations. However, single-cell analyses are essential for uncovering cell-to-cell variability and can powerfully support or contradict system-level models where traditional genetic perturbations face limitations. We present a single-cell method to quantify resection by integrating an optogenetic expression system at defined distances from an inducible double-strand DNA break (DSB) site in the budding yeast genome. Using this system, we test competing, unresolved models of how the DDC ‘counts’ DSBs and determines when to override the checkpoint. Current models propose that the extent of DNA damage is signaled by resection, either through liberated single-stranded DNA (ssDNA) or proteins bound along the non-resected strand. Although mechanistically plausible and widely known, these models rely on inconclusive evidence from gene knockout studies. An alternative hypothesis is that the DDC counts DSBs digitally, using factors located at 3’ break ends or at ss/dsDNA junctions. Here, we leverage natural cell-to-cell variability in resection rates as an intrinsic perturbation, avoiding the limitations of prior genetic approaches. Our single-cell data challenge models in which DNA damage is inferred from the extent or rate of resection or from proteins bound along resected DNA. To explore alternative mechanisms, we investigated DDC proteins localized at 3’ DSB ends or ss/dsDNA boundaries. By dynamically depleting candidate proteins after checkpoint arrest, we identified ss/dsDNA boundary proteins Ddc1, Dpb11, and Rad9 as essential for DDC maintenance and promising candidates for a cascade that acts as a digital DSB counter. Our findings demonstrate that quantitative, system-level single-cell approaches, coupled with dynamic perturbations, can resolve fundamental questions in DNA repair and checkpoint signaling.

## Introduction

DSBs pose a threat to genomic stability. If left unrepaired or repaired incorrectly, DSBs can lead to chromosomal rearrangements, mutations, or cell death, which are hallmarks of cancer and other genetic disorders^1–3^. Resection, the process degrading 5’ ends of broken DNA to generate 3’ single-stranded overhangs, is a pivotal step in DSB repair and influences the pathway choice between homologous recombination (HR) and non-homologous end joining (NHEJ)^4^. It enables HR by creating the ssDNA necessary for homology search and strand invasion and is thus critical to cellular integrity and preventing disease.

In budding yeast, which has long served as a eukaryotic model for elucidating DNA damage repair and checkpoint biology, a DSB is recognized by the MRX complex (Mre11-Rad50-Xrs2), which acts in conjunction with Sae2 to initiate end resection^5,6^. This limited, early resection enables displacement of bound proteins from the break such as the Ku complex (Yku70 and Yku80), needed for NHEJ, and trims back the DNA ends, creating short 3’ single-stranded overhangs^7–9^. Subsequently, more extensive resection is carried out by two main pathways: one involving Exo1, a 5’ to 3’ exonuclease, and the other driven by the helicase-nuclease complex Sgs1-Dna2^7,10^. These enzymes work to degrade the 5’ ends of the DNA strands, leaving long 3’ ssDNA tails. The ssDNA is coated by Replication Protein A (RPA), which serves as a platform for the recruitment and activation of the Mec1-Ddc2 kinase complex (homologous to ATR-ATRIP in humans)^11^. Mec1 phosphorylates the downstream effector kinase Rad53 (homolog of CHK2), with the assistance of mediator proteins such as Rad9^12,13^. Phosphorylated Rad53 amplifies the checkpoint signal and arrests the cell cycle pre-anaphase^14^.

Despite the presence of persistent DNA damage, budding yeast and mammalian cells eventually override or ‘leak’ through the DNA damage checkpoint arrest^15–19^. Override times strike an optimal balance between risk and speed in budding yeast and increase with the number of DSBs, from 6-10 hrs for one DSB, to 12-16 hrs for two DSBs^20^. (Previous work using another budding yeast strain also showed an increase in the arrest time with DSB number, albeit a more drastic jump from 8 hrs for one DSB to essentially ‘infinite’ for two DSBs^21^.) Mechanistically, increasing DSB numbers lead to stronger Rad53 hyperphosphorylation, which in turn maintains the DDC active for a longer period^22,23^.

Most models explaining how the number of DSBs tunes the strength of the DDC fundamentally center on resection: Liberated ssDNA oligonucleotides could stimulate Rad53 activity by binding the MRX complex^24^. In another resection-based model, the amount of exposed ssDNA determines the level of Mec1 activation, as larger regions of ssDNA allow for the recruitment of more Mec1-Ddc2 dimers, amplifying the checkpoint signal. This, in turn, could be expected to lead to the robust phosphorylation of downstream effectors such as Rad53, resulting in a stronger and more sustained cell cycle arrest^21^. There could also be priming and DNA synthesis processes competing with resection, with stretches of new dsDNA recruiting ss/dsDNA factors that amplify the DNA break signal. Again, the strength of the DSB signal would scale with the amount of resected DNA, which creates space for dsDNA islands. In contrast, there are less explored ways, independent of resection, by which cells could ‘count’ DNA breaks, for example, by factors located at the end of the 3’ non-resected strands or at ss/dsDNA junctions.

Although the resection-based DSB counting models are mechanistically plausible and were suggested by early genetic perturbation experiments, later work revealed contradictory or difficult-to-interpret results (Table 1). The original model was based on observations that two DSBs substantially delay override compared to one DSB, that is, the DDC counts DSB numbers. Further, in *yku70*Δ cells, resection is faster and most of the cells are permanently arrested. On the other hand, *mre11*Δ deletion leads to slower resection and suppresses the *yku70*Δ override defect, implying a positive correlation between resection rate and override time ^21^. This model was further supported by the observation that ATM (homolog of yeast Tel1, initiates checkpoint signaling before resection starts and Mec1 takes over^23^) activity is stimulated by ssDNA oligos in *Xenopus laevis* egg extracts and in human cells^24^. However, later experiments revealed that *fun30*Δ and *sgs1*Δ cells, which have slowed-down resection, were override incompetent^25^.

**Table 1:**
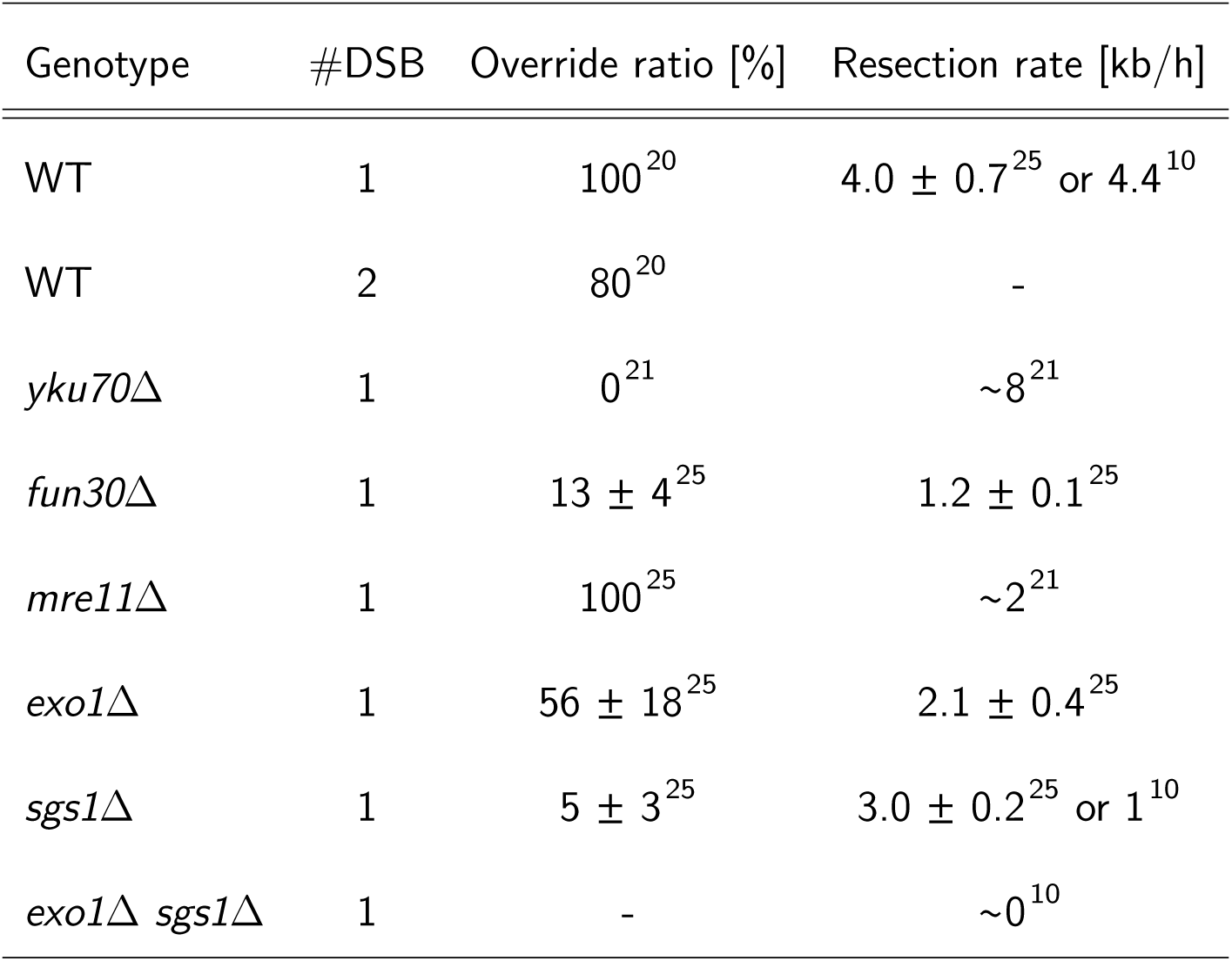
Previously reported resection rates and DDC override ratios in different genetic backgrounds. The override ratio is defined as the percentage of cells that overrode the DDC. ‘∼’ indicates that an approximate value was given in the publication.

Considering these difficult-to-reconcile results, we note that traditional genetic approaches face fundamental challenges in resolving this question: While the goal of genetic modification was to modulate resection rates and measure the resulting change in override timing, the genetic perturbation may actually affect both quantities independently, which would be difficult to disentangle. Furthermore, understanding the signal maintaining the DDC requires the checkpoint to be first initiated properly in order to study the subsequent override. However, permanent genetic perturbations may affect maintenance in addition to activation and establishment of the DDC.

Thus, our key idea was to pursue a different approach, namely, to leverage cell-to-cell variability, which represents a natural perturbation experiment of the resection rate. The DNA damage re-section, repair, and checkpoint machinery would remain entirely wild-type. If resection rates and override times would be observed to be correlated, which corresponds to analog counting of DSBs (Fig. 1 A), this would agree with resection rate-based models – although it would not be proof of a causal relationship. On the other hand, if these two quantities do not correlate (Fig 1 B), one could draw the stronger conclusion that resection cannot quantitatively determine the override time and thus the strength of the DDC. In this case, digital models of DSB counting would become plausible, to be supported by further evidence.

**Figure 1:**
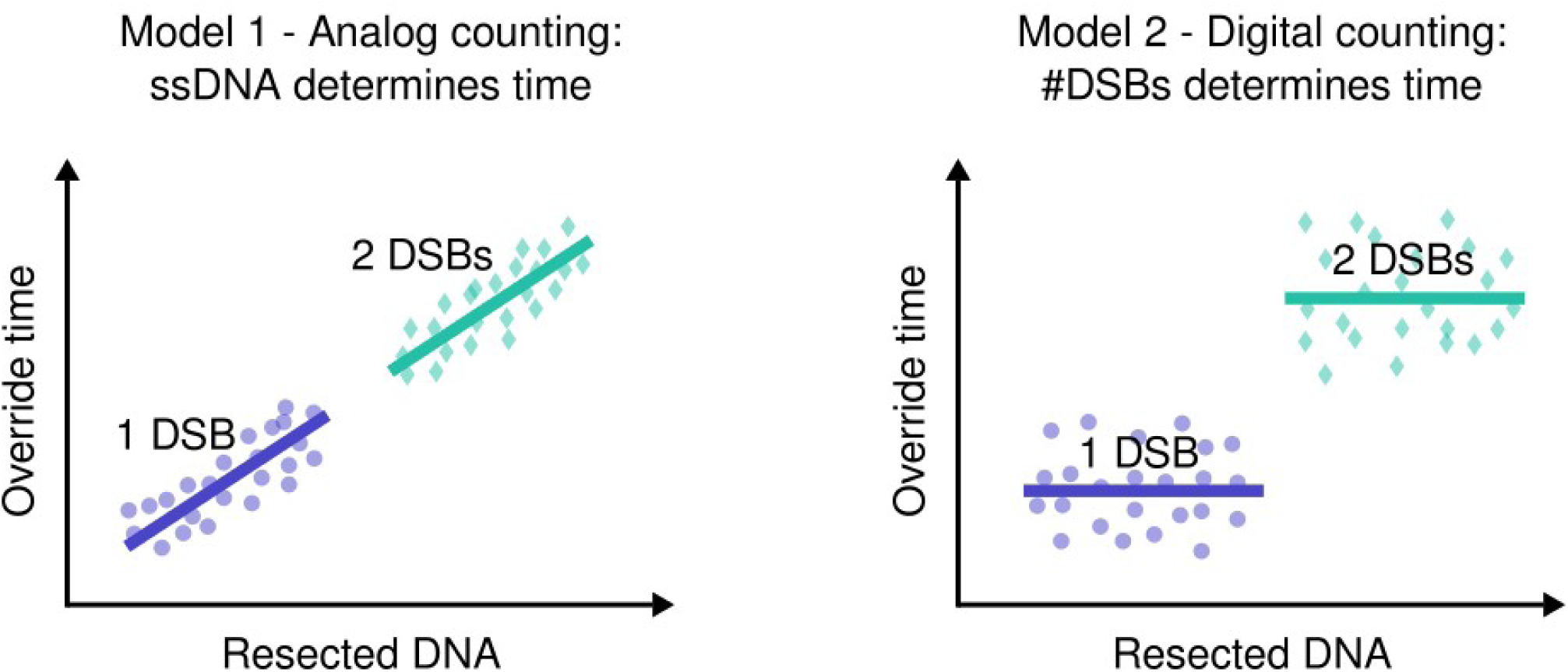
Possible outcomes of relating the resection rate to DDC override timing. A, B. Override time is correlated (A) or not (B) with the amount of resected DNA.

Resection has traditionally been measured in bulk cell culture using Southern blot analysis with strand-specific probes, which detect the progressive loss of dsDNA and the appearance of ssDNA ^26,27^. Quantitative PCR (qPCR) and next-generation sequencing (NGS) have also been used to map resection tracts and quantify the degradation of DNA near the break site^28,29^. At the sub-cellular level, the temporal and spatial organization of proteins involved in resection has been visualized with super-resolution microscopy^30^. Furthermore, a system was developed to quantify resection in fission yeast by monitoring the formation and disassembly of LacI-GFP and Rad52-mCherry fluorescent foci,^31^ which are challenging to identify and track reliably.

Here, we present:

- an optogentics-based tripwire system allowing resection to be quantified robustly at the single-cell level,
- characterization of the resection distributions indicating a faster median rate than is widely quoted in the literature,
- correlation of resection and override dynamics, which contradicts well-known resection-based models for DSB counting,
- evidence supporting that DSB counting occurs digitally by way of ss/dsDNA junctions.

We believe that the results of our novel approach to a long-standing question will advance the field and further spur the use of quantitative, single-cell approaches.

## Results

### Optogenetic tripwires

We realized that the processive loss of dsDNA and concomitant disruption of transcription ^32^ could be leveraged to quantify resection at the single-cell level. To this end, we developed a tripwire system consisting of a *light-inducible promoter* (*LIP*), controlled by the blue-light sensitive transcription factor El222, regulating the expression of the destabilized and fast-maturing fluorescent protein yEVenus-PEST^33,34^ (Fig. 2 A). The use of an inducible reporter generates a well-defined fluorescence peak when resection shuts off the expression of the reporter gene; in contrast, a constitutively expressed reporter requires distinguishing random fluctuations while expression is ‘on’ from the true end of the signal, which is more challenging. Furthermore, optogenetics avoids changes to metabolism and growth that arise when other inducible promoters such as the *GAL* or *MET3* promoters are turned on or off^35^. We introduced the optogenetic tripwire at various distances from an inducible DSB in previously validated^20,36^ budding yeast strains that could be arrested pre-start (in G1) before DNA replication by adding methionine to the growth medium (+M) and which could be released to start the cell cycle by removing methionine (-M)^37^. To create a DSB, we used a *GAL1pr-HO* construct to tightly control the activity of the endonuclease Ho. When switching to galactose medium (G), Ho creates a DSB at its recognition sequence (Ho cut site, *HOcs*) which was placed strategically either in the promoter of *MIC60* on chromosome XI and/or at the *URA3* locus on chromosome V. *MIC60* is located in a long genomic region that is not essential^38^, thus ensuring that the DSB does not result in death due to resection of essential genes. We used the cut site at *MIC60* for most of the experiments and, when needed for a second DSB, at *URA3*. Furthermore, the endogenous Ho cut-site at the mating type locus (*MATα*) was abolished by silent mutations in the *alpha1* gene (*MATα-syn*).

**Figure 2:**
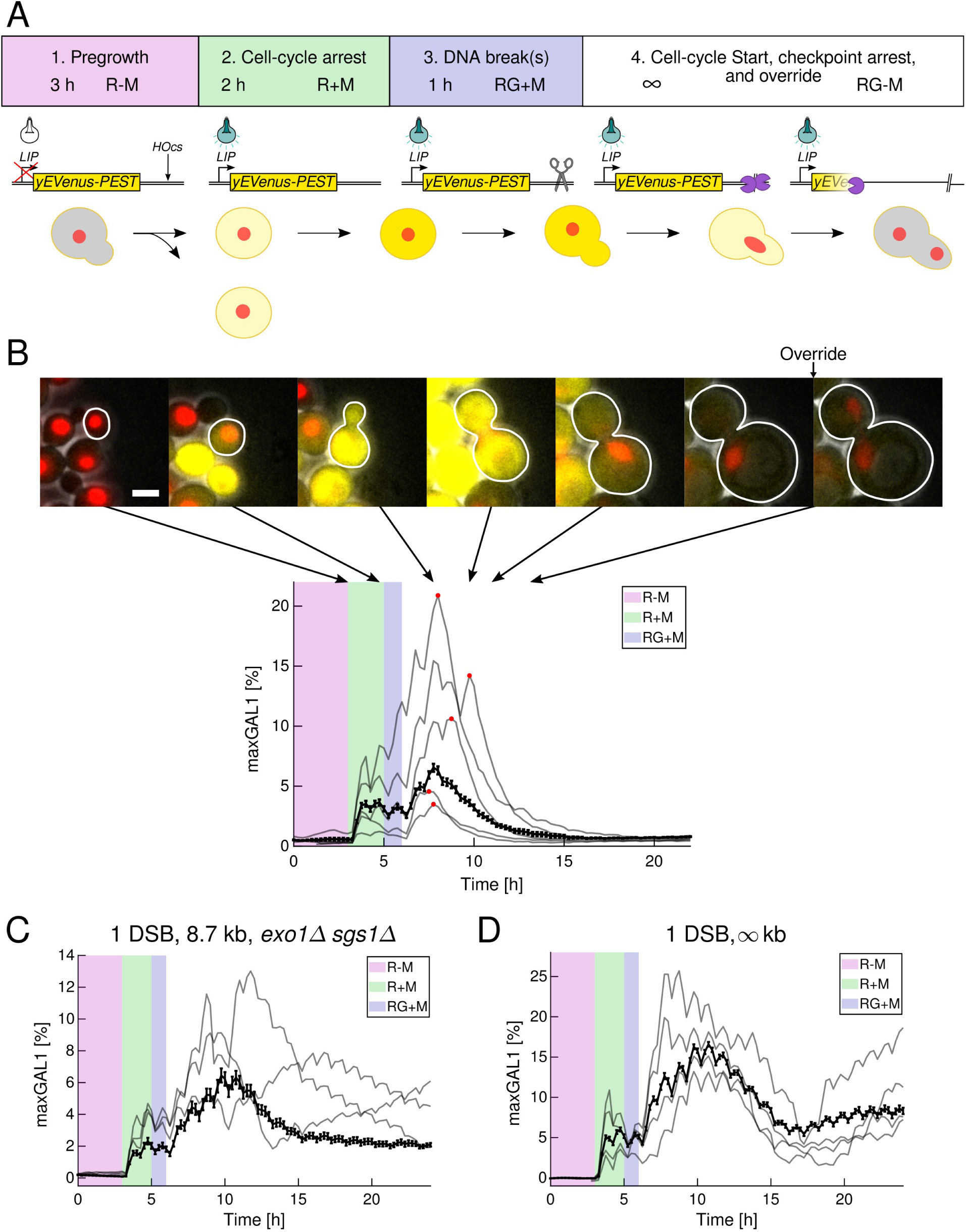
Optogenetic tripwire experiments. A: Experimental protocol. B. Representative images and fluorescence time courses of cells with the tripwire placed 8.7 kb from a single Ho cut-site in the *MIC60* promoter. The last two snapshots are one frame apart (15 min). The override time is defined as the interval between the time of budding and of anaphase, shown in the last image. Mean (bold trace) with standard error of the mean (error bars) and five representative fluorescence time courses of yEVenus-PEST. Red circles on the time courses indicate peaks which are interpreted as the time points at which resection eliminated the tripwire locus (**Methods**). The resection time is the difference in the time of this peak and of DSB induction, that is, the switch between R+M and RG+M at time point 5 h in this plot. Scale bar, 5 *µ*m. C. Mean (bold trace) with standard error of the mean (error bars) and three representative fluorescence time courses of *exo1*Δ *sgs1*Δ cells with the tripwire placed 8.7 kb from the *HOcs* locus. D. Mean (bold trace) with standard error of the mean (error bars) and four representative fluorescence time courses of cells with the tripwire placed on another chromosome as the *HOcs* locus. For all panels, the number of cells as well as the fraction of cells overriding are supplied in Supplementary Table 1.

Tripwire experiments followed a fixed protocol (Fig. 2 A): Initially, cells were grown in synthetic complete medium with raffinose and without methionine (R-M). They were then arrested in G1 for 2 hrs by addition of methionine (R+M), ensuring that all chromosomes were present as a single copy when cut in order to prevent repair by HR. Galactose was then added for 1 hr (RG+M), generating an Ho-induced DSB. Finally, cells were released from the G1 arrest and started a new cell cycle by removing methionine from the medium (RG-M). DSB repair was prevented by continuous induction of *GAL1pr-HO* in galactose medium. In addition to the media switches, cells were illuminated with pulsing light for 15 min every hour, starting at the time point when methionine was added until the end of the recording. The pulsing light regime allowed the induction of the optogentic tripwire while limiting phototoxicity^34^.

The tripwire typically produced a clear fluorescence peak (Fig. 2 B) whose timing differed from cell to cell and systematically with distance from the DSB. To validate our system, we placed the optogenetic tripwire in two different strains: In *exo1*Δ *sgs1*Δ cells (Fig. 2 C), resection is expected to be severely impaired with only 5-10% of cells resecting further than 3 kb from the DSB^7,10^. In our second control, the cut site was placed on another chromosome (*URA3* locus) than the tripwire (Fig. 2 D). In both strains, fluorescence increased as expected but peaked on average at 10.8 ± 2.0 h and 10.4 ± 1.1 h (mean ± STD) in *exo1*Δ *sgs1*Δ and the infinite-distance control, respectively. Thereafter, fluorescence slowly decreased, potentially due to a global reduction in transcription^39,40^. Despite the overall decline in fluorescence in most cells, fluorescence remained high in some cells. Yet, despite the decline, fluorescence did not go to zero in these controls, in contrast to the case where the decrease was caused by resection (Fig. 2 B). The controls also showed clear oscillations with the period of the light pulses, 1 hr, in addition to the overall rise and fall. We thus defined a set rules to identify the resection-induced peak computationally (**Methods**) (typical examples shown in Fig. 2 B). Note that it was key that the reporter system could be pulsed periodically for identifying the resection time (without affecting cell metabolism), as the light-induced oscillations showed the tripwire to be intact, while transcriptional output increased and decreased during the DDC arrest, making the peak over the entire time course an unreliable indicator.

### Noise in initiation and speed of resection

To characterize resection rate at the single-cell level, we placed the tripwire at various distances from the *HOcs* locus in the *MIC60* promoter (Fig. 3 A, B). Interestingly, as time progressed, the distribution of resection times flattened only in the faster half of the population (5^th^ and 25^th^ percentile regression lines, Fig. 3 B); in this sub-population, the resection rate varied markedly between cells. On the other hand, the slower half of the population (right of the median) showed a remarkably stereotyped resection rate; here, cell-to-cell variability stemmed from the initiation time of resection. The source of the initiation time variability is not simply noise in cell cycle Start since budding and resection time are not significantly correlated (Supplementary Fig. 1). (At the timescale of interest, budding is a reliable indicator of cell cycle Start as it follows Whi5 nuclear export by 15 ± 8 min (mean ± STD)^41^.) Instead, the data indicate that initiation of resection follows DSB induction by 84 ± 6 min (mean ± STD) (preceding budding by about 30 min), with a long delay of up to 130 min in half of the population.

**Figure 3:**
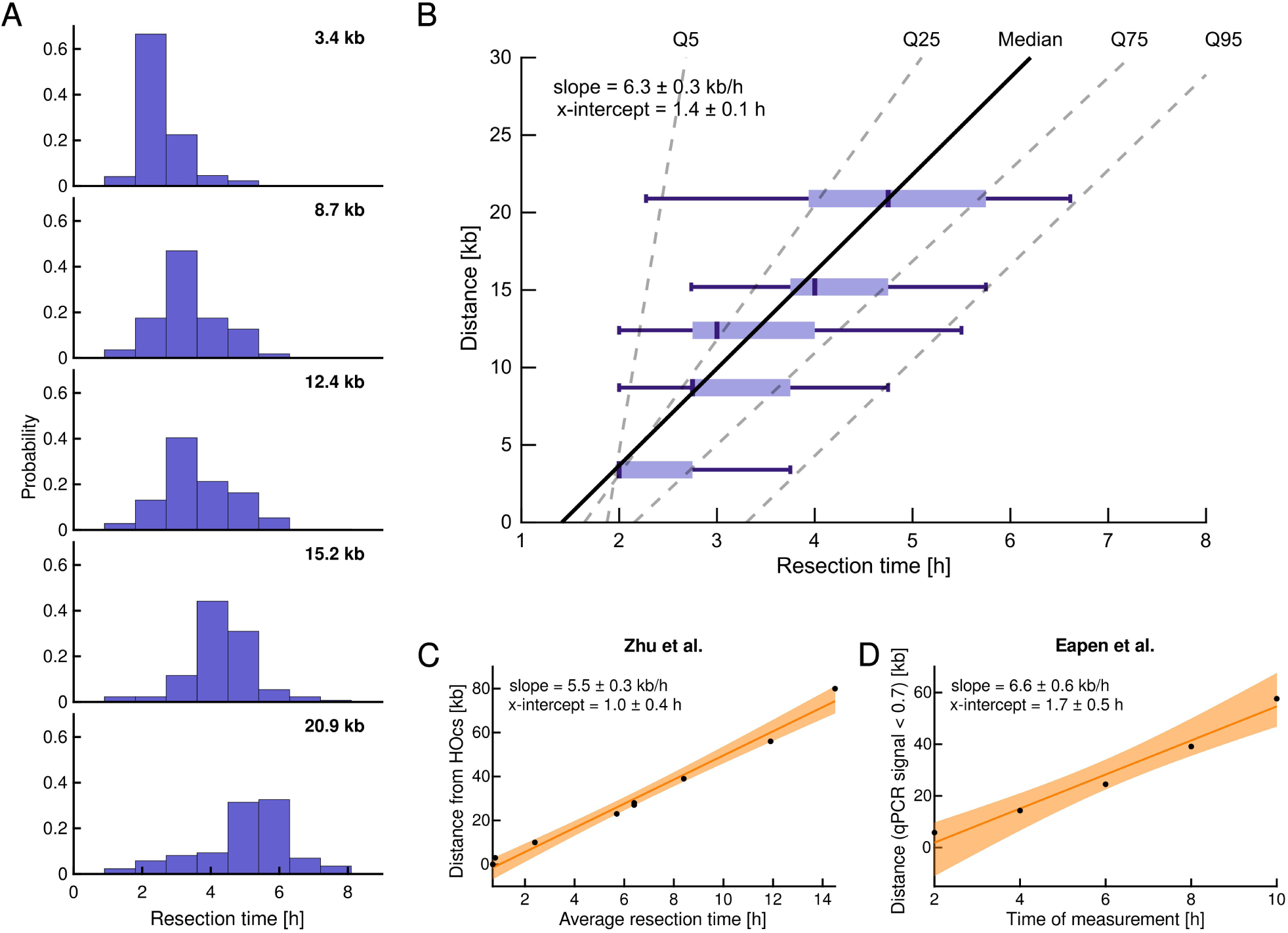
Resection initiation, rate, and distribution. A. Histograms of the resection time for different tripwire distances. B. Distributions of resection times. Boxes indicate 25^th^ and 75^th^ percentiles. Dark, vertical lines inside the box indicate medians of the respective distributions. Whiskers delineate the 5^th^ and 95^th^ percentiles. The oblique black line represents a linear fit to the medians, whose slope and x-intercept are indicated with the standard deviation, computed by bootstrapping (**Methods**). The four dashed gray lines are linear regressions on the 5^th^, 25^th^, 50^th^, 75^th^, and 95^th^ percentiles. C: Plot of the values in Zhu et al. ^10^ (circles) and linear regression. D. Plot of the values in Eapen et al. ^25^ (circles) and linear regression. In C and D, the shaded area is the 95% confidence interval of the fit. In B, C and D, all the linear regressions are computed on the horizontal residuals, as the distance is the independent variable. The dependent variable, that time, is on the x-axis, so that the slopes correspond to the resection rates. For all panels, the number of cells as well as the fraction of overriding cells are provided in Supplementary Table 1.

By performing a linear regression analysis of the median resection time at different distances, we determined the resection rate to be 6.3 ± 0.3 kb/h (mean ± STD) (Fig. 3 B), which represents a large deviation from the widely reported value of ∼4 kb/h^10,25,28,42^ To elucidate this divergence, we re-evaluated results from Zhu et al.^10^ and Eapen et al.^25^ by linear regression analysis (Fig. 3 C, D). In both, a DSB was produced by turning *GAL1pr-HO* on in a cycling budding yeast population. Zhu et al.^10^ computed resection rates in bulk populations by dividing the resected length of DNA by the time of resection, using the switch to galactose medium as time point zero. Because resection is influenced by cell cycle stage and requires specific steps to initialize (including DNA nicking by the MRX complex and removal of the Ku complex^8,9^), we propose to infer the zero time point from the data by linear regression, which then yields 5.5 ± 0.3 kb/h (slope ± SE) for the resection rate based on the same data (Fig. 3 C). Eapen et al.^25^ performed linear regression but only for earlier time points (before 6 hrs) and excluded later time points. Considering all time points, the resection rate becomes 6.6 ± 0.6 kb/h (slope ± SE) (Fig. 3 D). In conclusion, based on our careful measurements – involving synchronized cell cycle Start and single-cell resolution – we propose to revise the widely quoted resection rate to 6.3 ± 0.3 kb/h (mean ± STD) and to consider a delay of 84 ± 6 min (mean ± STD) with respect to DSB induction.

### Mutations and number of DSBs affecting resection

We analyzed the resection rates of several mutant strains or in 2-DSB cells using the same approach but only two or three tripwires (Fig. 4 A-D). The first two deletions affected resection directly: Long-range resection following a DSB is performed by the exonuclease Exo1 and by the endonuclease Dna2 in complex with the helicase Sgs1^7,10^. To compare our measurements with previous studies, we divided the mutant resection rates by the wild-type rate (Fig. 4 E). Our results were highly comparable for *exo1*Δ and *sgs1*Δ and qualitatively similar for *fun30*Δ. Next, we measured the resection rate around one break in the presence of a second break, which, to the best of our knowledge, has not been quantified previously. Thus, we added a second *HOcs* on another chromosome (at the *URA3* locus) and used three tripwires. Interestingly, the presence of a second DSB substantially reduced the resection rate compared to 1 DSB (by ≈35%).

**Figure 4:**
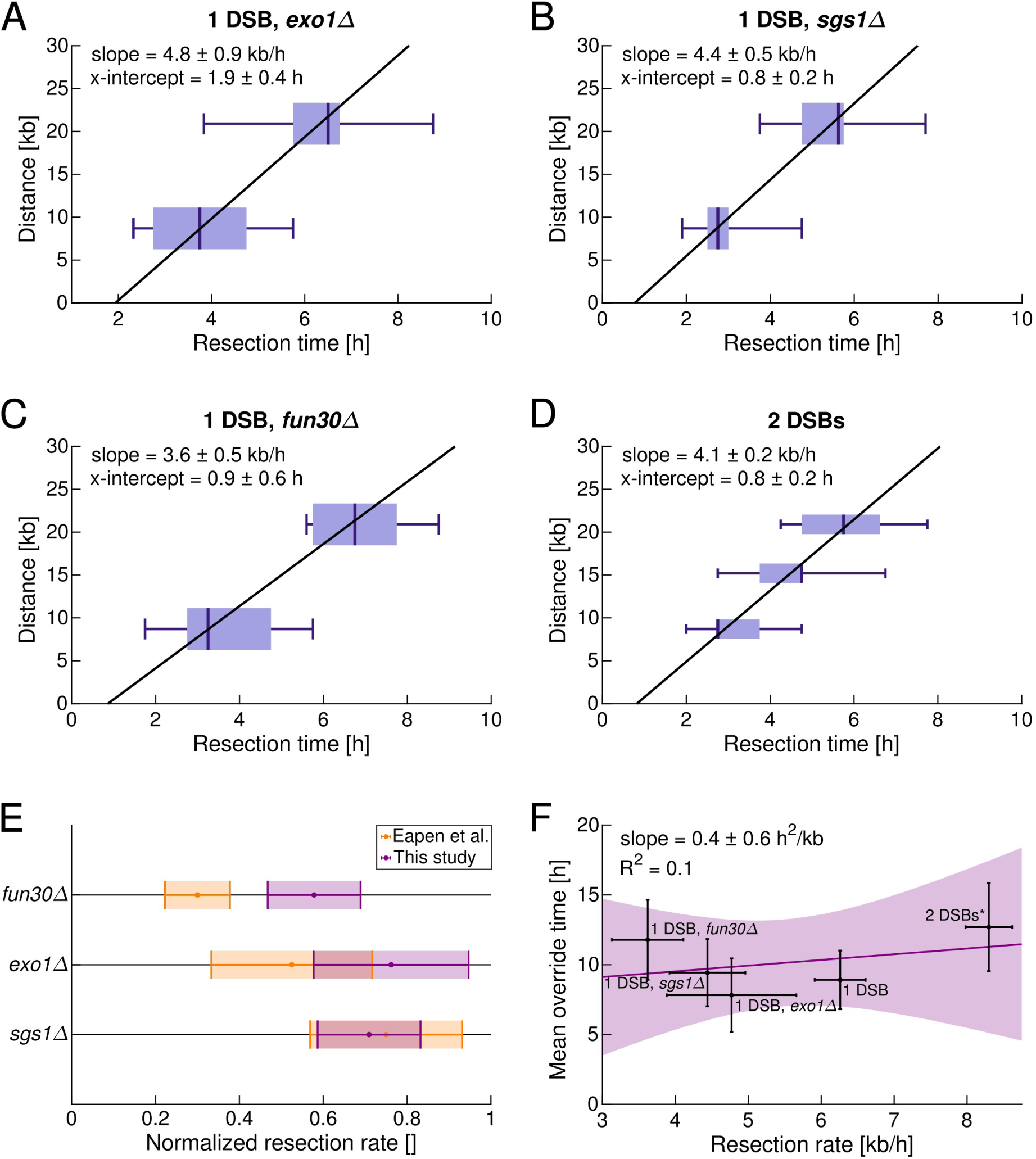
Effects of resection-related genes as well as a second DSB on resection. A, B, C, D. Distributions of resection times in *exo1*Δ (A), *sgs1*Δ (B), *fun30*Δ (C), or 2-DSB (D) cells. E. Comparison between our results and Eapen et al. ^25^ (Table 1). To account for the discrepancy in the measurement methods, the resection rate was normalized with respect to the respective wild-type results. Error bars, STD. F. Mean override time as a function of the resection rate, defined by the slopes of the linear regressions in panels A, B, C, D and Fig. 3 B. Override times were pooled from all tripwire experiments in each panel A, B, C, D, and Fig. 3 B. ‘2 DSBs*’ indicates that the resection rate in the 2-DSB cells was doubled to estimate the amount of resected DNA. For all panels, the number of cells as well as the override ratios are provided in Supplementary Table 1.

Finally, we correlated these resection rate data with the additionally measured override times scored for each cell in each strain. We observed no significant correlation (Fig. 4 F). Thus, these results do not support the resection-based model for counting DSBs. Note that the resection rate serves as a proxy for the amount of resected DNA; consequently, the ‘total resection rate’ in the 2-DSB cells is doubled in Fig. 4 F.

### Single-cell-based correlations discriminate DSB counting models

To more fully leverage our methodology and to avoid uncertainties inherent in interpreting the effects of gene deletions (Fig. 4 E), we focused on cell-to-cell variability and analyzed the correlation between override time and resection at the single-cell level (Fig. 5). We tested tripwires at three different distances (8.7 kb, 15.2 kb, and 20.9 kb) to allow for different contributions to cell-to-cell variability from resection initiation and resection speed: At the shortest distance, override initiation variability should dominate; at the longest distance, variability in resection speed should play a substantial role (Fig. 3 B). To obtain a single-cell resection rate, we set the initiation time of resection to the average values for 1-DSB cells (Fig. 3 B) or 2-DSB cells (Fig. 4 D), respectively. For shorter resection times, that is, faster resection rates, the data became sparse due to the 15-min imaging intervals. Thus, we excluded cells with resection rates faster than 12 kb/h. The full data set is also presented in terms of the raw resection times (roughly, the inverse of resection speed) in Supplementary Fig. 2 and support the same conclusions.

**Figure 5:**
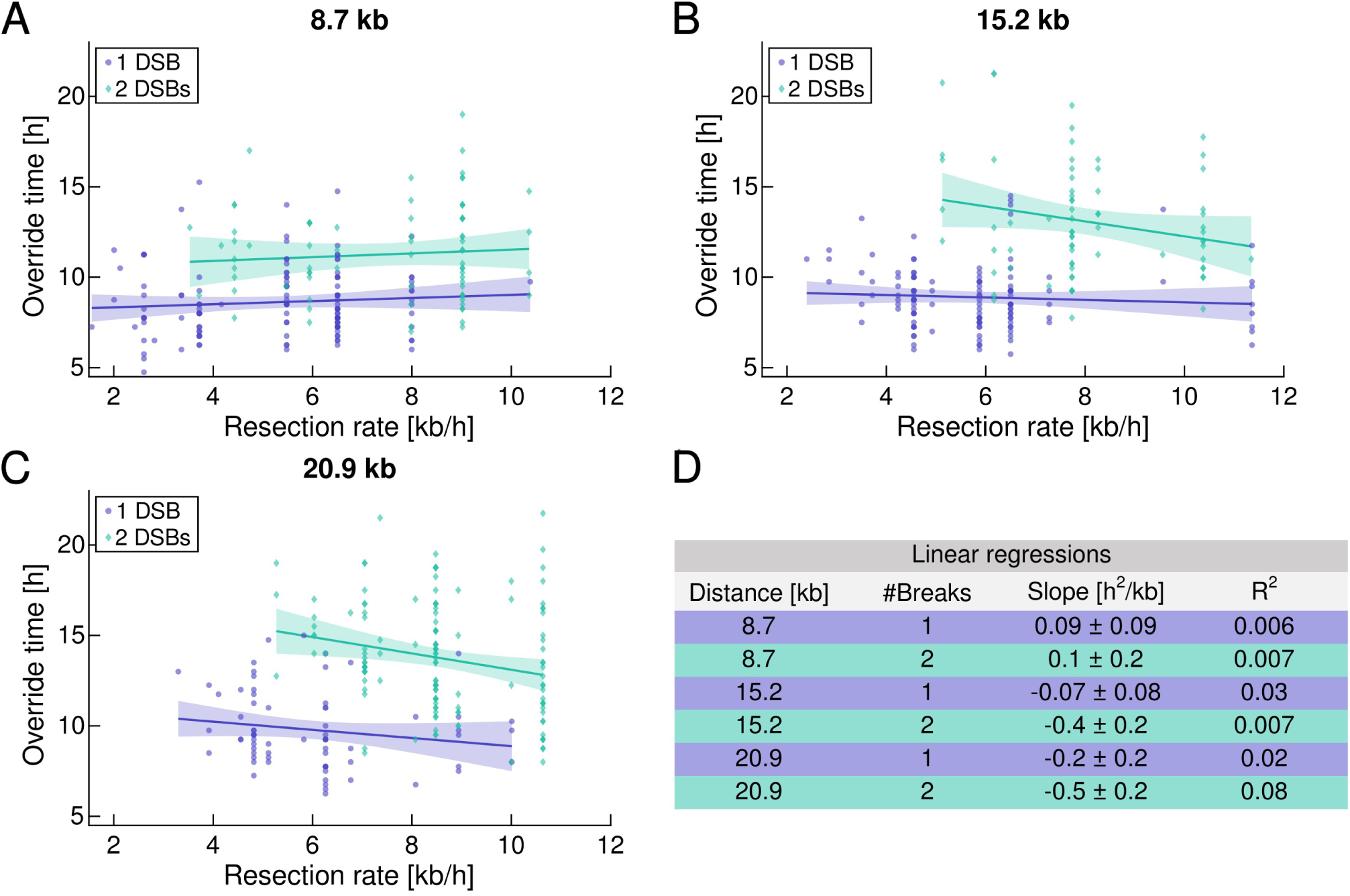
No correlation or weakly anti-correlated relationship between override time and resection rate. A, B, C. Override time as a function of resection rate in 1-or 2-DSB cells with the tripwire placed at 8.7 kb (A), 15.2 kb (B) and 20.9 kb (C) from an Ho cut-site in the *MIC60* promoter. For cells with 2 DSBs, the resection rate was doubled to estimate the total amount of resected DNA per unit time. Best linear regression line shown with the 95% confidence interval shaded. D: Characteristics of the linear regressions. For all panels, the number of cells as well as the override ratios are supplied in Supplementary Table 1.

The correlations between override times and resection speeds were poor (all *R*^2^ ≤ 0.08), contradicting models where resection leads to strengthened checkpoint arrest (Fig. 5). Specifically, for the 8.7 kb tripwire, the regression lines for both 1- and 2-DSB cells were essentially flat. Curiously, at 15.2 kb and 20.9 kb, a mildly *negative* linear regression line emerged, particularly in the presence of two DSBs, associating faster resection with a *weaker* checkpoint. While the absence of a positive correlation contradicts resection-based DSB counting, the presence of a (mildly negative) correla-tion does not necessarily imply a causal relationship between resection and checkpoint strength but shared upstream regulation.

### ss/dsDNA junction proteins maintain the DDC and could count DSBs

To identify alternative DSB counting mechanisms, we used gene deletions to examine candidate DNA break response proteins that are known to be localized near break ends or at the ss/dsDNA boundary (Fig. 6 A). An early initiator of the DNA damage response, Mre11 is a core component of the MRX complex. Mre11 plays a central role in DSB recognition and in initial nicking of DNA prior to long-range resection, and contributes to DSB end tethering^43^. *mre11*Δ deletion led to a significant reduction in override times in the presence of one or two DSBs. Given that Sae2 enhances MRX’s nuclease activity^6^, we deleted *SAE2* and observed that the override time was indistinguishable from wild type, suggesting that MRX nuclease activity is not responsible for *MRE11*’s role in maintaining wild-type override timing. MRX complex formation is followed, among other processes, by long-range resection. In agreement with the work already presented, *exo1*Δ *sgs1*Δ deletions, which essentially prevent resection^7,10^, only mildly reduced override times compared to wild type. Next, we considered the Dpb11 scaffold, which is recruited to ss/dsDNA junctions by the 9-1-1 clamp complex, which in turn incorporates Ddc1^44,45^. Dpb11 and Ddc1 both harbor Mec1-activating domains and contribute to Rad9 recruitment to the break^46,47^, with Dpb11 binding simultaneously Ddc1 and Rad9^48^. Mec1-mediated phosphorylation of Rad9 promotes oligomerization and binding of Rad53^12,49^, which in turn can be activated by Mec1^50^. Due to *DPB11* essentiality, we only deleted *DDC1* and *RAD9*. Both *ddc1*Δ and *rad9*Δ deletions caused extremely short budding-to-anaphase times in the presence of one or two DSBs. Based on their localization, Mre11, Dpb11, and Ddc1 could represent candidates for digital DSB counting; however, the effects of their deletions could also be explained by roles in initial establishment of the DDC.

**Figure 6:**
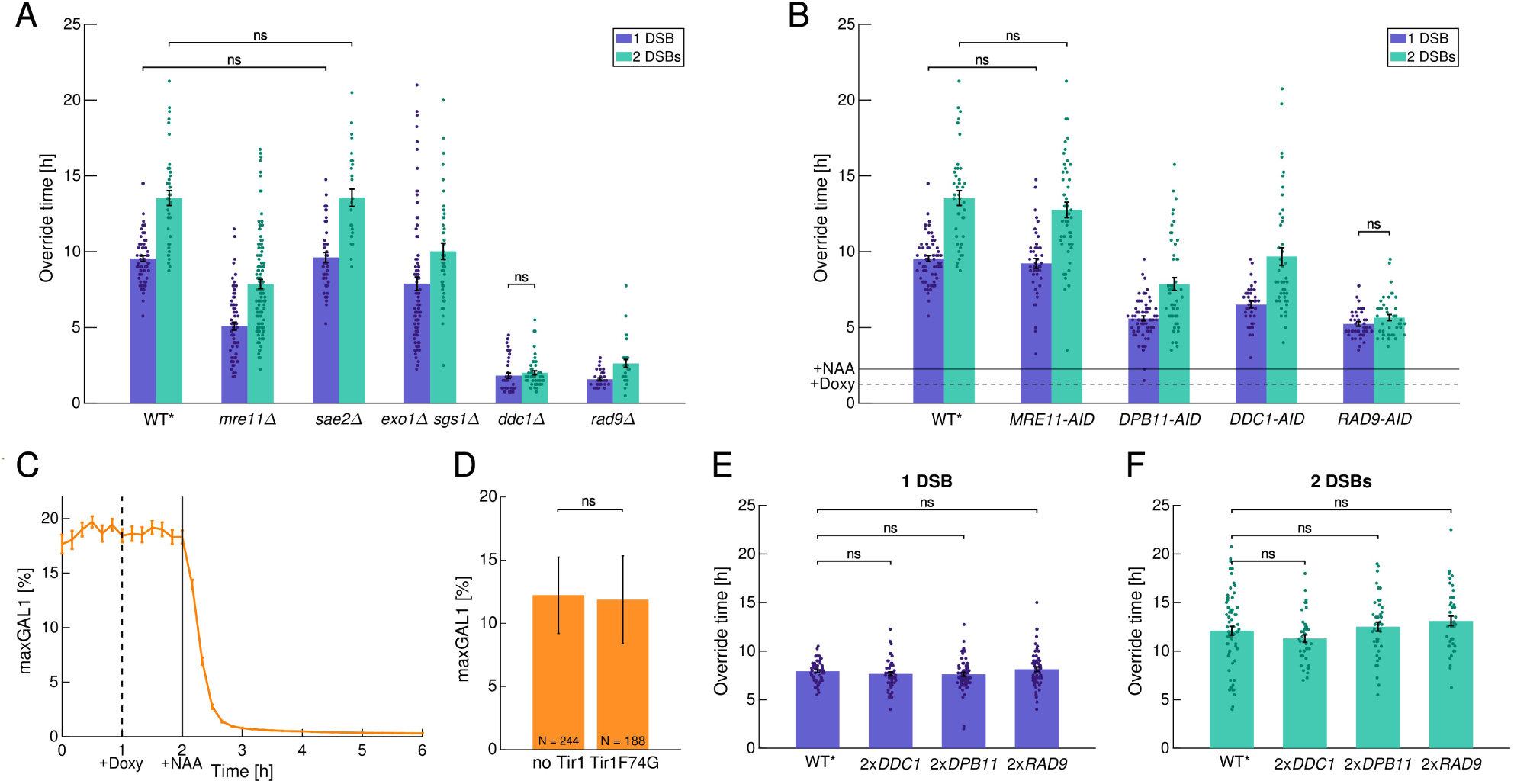
Deletion, induced degradation, and over-expression analyses suggest that ss/dsDNA junction proteins enable DSB counting. A, B. Mean and standard error of the mean of the override time for cells with 1 or 2 DSBs and specific DDC genes deleted (A) or knocked down post arrest (B). C, D. Validation of AID depletion system using Tir1(F74G) and NAA. C. Cells were grown in synthetic complete with 2% glucose, without methionine (D-M). Doxycycline and NAA added at 1 h and 2 h respectively, following the same timing as in B. D. To evaluate the leakiness of the system, cells with or without Tir1(F74G) were grown in D-M and imaged on glass slides. N = 87 at 2 h. E, F. Mean and standard error of the mean of override times for strains with 1 DSB (E) or 2 DSBs (F) and a second copy of the indicated DDC genes. A, B, E, F. WT* indicates strains with the same genetic background as the other strains in the respective panel but without the indicated mutations or second gene copies. A, B, D, E, F. Only relationships, which were not significant different by two-sample Student’s t-test analysis are highlighted. For panels A, B, E, F the numbers of cells as well as the override ratios are provided in Supplementary Table 1.

In order to specifically identify roles in maintaining DDC arrest, we switched to dynamically depleting DDC proteins after arrest (Fig. 6 B). We employed an auxin-inducible degron (AID) system and tagged individual proteins of interest with the AID* tag^51,52^. To reduce leaky degradation, we employed the Tir1(F74G) enzyme variant^53^, and additionally controlled expression of *TIR1(F74G)* with the doxycline-inducible Tet-On system^34^. We induced protein degradation by adding doxycycline at 2 h after methionine release (approximately 75 min post budding) and added naphthalene-acetic acid (NAA) one hour thereafter. NAA produces less phototoxicity in combination with fluorescence microscopy compared to the more commonly used indole acetic acid (IAA)^54^. To test our system, we used a reporter strain with *MET3pr-AID*-ymScarletI* and observed fast degradation and low leakiness (Fig. 6 C, D).

We first depleted Mre11 after checkpoint activation. In contrast to *mre11*Δ deletion, we observed no significant differences in override timing, suggesting that other Mre11 functions besides nuclease activity, e.g., tethering of broken ends, do not set the strength of the DDC either. The primary effect of the *mre11*Δ deletion may be in perturbing establishment of DDC arrest. Next, we examined the roles of ss/dsDNA proteins by knock-down after checkpoint activation. Indeed, degradation of Ddc1, Dpb11, and Rad9 resulted in similar, substantial reductions in override time. Given that Ddc1 and Dpb11 act upstream of Rad9, their effect on DDC strength may be through Rad9 phosphorylation and Rad53 activation. All three proteins, associated with ss/dsDNA junctions^47,55–60^, are essential for maintaining DDC arrest, suggesting a straightforward mechanism for DSB counting.

For this model to be realistic, it is important that only Ddc1, Dpb1, or Rad9 proteins associated with ss/dsDNA junctions contribute to DDC maintenance, not potentially unbound protein in the nucleus or the cytoplasm. To test this requirement, we doubled the copy numbers of *DDC1*, *DPB11*, and *RAD9* genes, increasing the amount of each protein globally by about two fold^61^, especially affecting any potential pool of protein that is unbound but signaling. Both in the presence of one (Fig. 6 E) and two DSBs (Fig. 6 F), the second gene copies had no statistically significant effect on override timing. In conclusion, knock-downs showed that all three proteins associated with ss/dsDNA junctions are essential for maintaining DDC arrest but increasing their concentrations globally has no effect. Thus, ss/dsDNA junctions are promising digital DSB counters, plausibly through well-established Rad9-mediated phosphorylation of Rad53.

## Discussion

DDC arrest varies with the number of DSBs^20,21^. To elucidate the mechanisms of DSB counting, i.e., how increasing the number of DSBs strengthens DDC arrest and delays override, previous work relied on constitutive mutations that led to difficult-to-reconcile results. We developed an optogenetic tripwire system that tested whether the amount or rate of resected ssDNA could be the source of the DSB count signal. For this, we leveraged the natural cell-to-cell variability in the override process and found no correlations (or weakly negative correlations) at the single-cell level between the override time and resection rate.

Furthermore, we observed that once resection was initiated, elongation proceeded at a rate of 6.3 kb/h. This rate is substantially faster than the widely reported ≈4 kb/h resection rate^10,25,28,42^. The previous estimate did not distinguish between the actual resection speed and the time to initiate resection. By aligning resection timing across multiple genomic distances, we disentangled these processes and found an average delay of about 1.4 h between DSB induction and the start of resection. Importantly, resection timing was not simply determined by cell-cycle Start (G1/S transition) since the timing of budding showed no significant correlation with resection onset. Instead, the delay potentially reflects the time required for early processing events at the DSB such as Ku removal and initial DNA nicking by MRX. Our re-analysis of previous data supports this two-step model: When allowing a delayed start, published resection data^10,25^ yielded regression slopes of ≈5.5-6.6 kb/h, consistent with our single-cell-based rate.

Together, our data suggest that cells count DSBs in a digital manner, setting the DDC strength and override time based on signals from the ss/dsDNA boundary, rather than based on an analog signal from resected ssDNA. To uncover the molecular basis of this digital counting mechanism, we performed genetic deletions, dynamic protein depletion, and mild over-expression in cells with one or two DSBs. This approach aimed to identify early DNA damage response factors which are needed for DDC arrest maintenance, which bind limited partners such as ss/dsDNA junctions, and whose impairment could eliminate the difference in checkpoint duration between single-DSB and double-DSB cells, such as *RAD9*.

Our results further indicate that cells distinguish multiple breaks based on processes downstream of MRX activation since Mre11 depletion did not affect DDC maintenance. Similarly, the *exo1*Δ and *sgs1*Δ deletions only slightly shortened the DDC arrest. Combined with the single-cell resection speed measurements, these findings indicate that neither the amount of liberated ssDNA nor proteins bound along resected ssDNA count DSBs. On the other hand, disrupting factors that localize to the ss/dsDNA junction had a pronounced effect, suggesting that the ss/dsDNA boundary may be the source of DSB counting. Ddc1, a subunit of the 9-1-1 checkpoint clamp loaded at ss/dsDNA^55,56^; the scaffold protein Dpb11, which links the 9-1-1 clamp to Mec1 and Rad9^47,58,60^; and Rad9, a checkpoint mediator that binds chromatin around damage sites^46,47,59,60^, proved to be potential candidates for digital DSB counting.

Beyond specific mechanistic insights, our work highlights the broader value of quantitative, dynamic single-cell approaches in DNA repair and checkpoint biology, more commonly applied in other areas of quantitative biology^62,63^. Leveraging cell-to-cell variability allowed circumventing the genetic perturbations, which can have pleiotropic effects. Dynamic perturbations allowed focusing on check-point maintenance. Continued development of such approaches is likely to produce powerful other approaches needed to elucidate challenging mechanistic questions in the field.

## Methods

### Plasmid construction

All plasmids were constructed and propagated using *E. coli* DH5*α* or XL10. Plasmids were constructed either by Gibson assembly (NEB, USA) or by restriction-enzyme-based cloning. DNA digestion and ligation were performed using restriction endonucleases and T4 DNA ligase from NEB, USA. All PCRs were performed with Phusion Polymerase (NEB, USA). For growing bacteria, we used LB medium supplemented with ampicillin. All constructs were verified by Sanger sequencing (Microsynth AG, Switzerland). The list of plasmids used is provided in Supplementary Table 2.

### Strain construction

All experiments were performed with budding yeast strains with the W303 background (*ade2-1 leu2-3 ura3-1 trp1-1 his3-11,15 can1-100*). Transformations were performed using the standard lithium acetate method^64^, and transformed strains were selected using dropout plates or plates containing antibiotics. Crosses were performed by standard methods for budding yeast mating, sporulation, tetrad dissection, and tetrad selection. Plasmid integration and construct activity were verified by fluorescence microscopy after induction of the constructs. Strains that showed fluorescence were screened for single-copy integration using PCR with primer sets that allowed one or several copies of the construct in the genome to be distinguished. To avoid inserting extraneous DNA that could affect DNA repair processes or the checkpoint, we repeatedly used the pop-in/pop-out method^65^ to leave no marker in the genome whenever possible. The basic genotype for all strains was *cln1*Δ *cln2*Δ*::MET3pr-CLN2* (promoter replacement) *cln3*Δ*::GAL1pr-HO HTB2-mCherry::HIS5 MATα-syn*. In addition, the *rad5-G535R* mutation of W303 was corrected^66^. HOcs stands for a sequence of 30 base pairs, namely, TTCAGCTTTCCGCAACAGTATAATTTTATA. The tripwire is placed in such a way that the non-coding strand is resected. To add a second copy of a gene, we selected 1000 bp for the promoter and 300 bp for the terminator. If another gene overlapped with this selection, we stopped the promoter/terminator before this next gene’s open reading frame.

For knock-down experiments, we used the residues of AID* defined in Morawska et al.^51^ but with the yeast-optimized nucleotide sequence from Lu et al. ^52^. To prevent leaky degradation, following Lu et al.^52^, we controlled the expression of *Tir1(F74G)* ^53^ with the Tet-On system^34^. The list of budding yeast strains used is provided in Supplementary Table 3.

### Media and growth conditions

We used synthetic complete media (SC) supplemented with 2% (w/v) glucose (D), 3% (w/v) galactose (G) or 3% (w/v) raffinose (R). To arrest the *cln1*Δ*, cln2*Δ*::MET3pr-CLN2 cln3*Δ*::GAL1pr-HO* cells in G1 phase or release them from G1 arrest, methionine was added (+M) or removed from the medium (-M), respectively. All experiments followed a fixed protocol: pregrowth in R-M, 2 h in R+M, 1 h in RG+M, and switch to RG-M. For AID system experiments, we added 10 *µ*M doxycycline (Sigma-Aldrich Chemie GmbH, Switzerland) after 2 h of RG-M and added 1 mM NAA (Sigma-Aldrich Chemie GmbH, Switzerland) 1 h after doxycycline. For microscopy experiments, cells were grown in CellASIC ONIX microfluidic plates for haploid yeast cells in media controlled by the ONIX2 microfluidics system (Merck, Germany). For routine growing of yeast we used solid SCD plates with 2% agar.

### Microscopy and optogenetics

Time-lapse recordings were carried out with a Plan Apo *λ* 60x/1.40 oil objective and a Hamamatsu ORCA-Flash4.0 camera. The microscope was operated using Nikon NIS Elements AR 5.21.03 64-bit software. To decrease the potential phototoxicity of imaging during long arrests, the interval between the images was set to 15 min. For imaging, the exposure time of all channels (brightfield, RFP and YFP) was 100ms. To optogenetically induce fluorescent reporters, we used the ‘white’ diascopic LED light source, as in previous work^34,67^.

### Image analysis

For segmenting the microscopy images, we used the YeaZ convolutional neural network and graphical user interface^68^. The override time was manually scored as the difference between the time point at which a cell with damaged DNA budded and the time point at which the Htb2-marked nuclei separated.

### Data processing

We excluded from our data set:

- cells which are budded at the time of DSB induction,
- cells which do not immediately re-arrest at G2/M after cell cycle release in RG-M medium,
- cells which have unusual behaviors such as rebudding before division or endomitosis.

The resection time is defined as the difference between the peak time and the break induction time (switch from R+M to RG+M). To detect the peak and tripwire extension by resection, we selected the latest peak with a prominence greater than a 5% of the maximum of the time course, using the ‘findpeaks’ function in Matlab. Selecting the latest peak and not the highest allows to account for tripwire fluorescence which reaches a noisy plateau. We limit the minimal prominence to prevent findpeaks’ from detecting the oscillations due to the 1-hour period of the light induction protocol. All traces and peak calls were checked manually and misidentifications corrected.

The distance between the yEVenus stop codon and the *HOcs* locus defines the tripwire distance.

### Reproduction from the literature

In Fig. 3 C, we plotted the values given in the table of Fig. 1 C of Zhu et al.^10^. For Fig. 3 D, we reproduced the 1-DSB resection data from Eapen et al.^25^ given in Fig. 3 D, E, F.

### Statistics and reproducibility

No statistical method was used to predetermine the sample size.

## Data Availability

All data supporting the findings of the study are available in the article and Supplementary Information.

## Acknowledgments

We thank Roxane Dervey for help with strain construction and media preparation; Ara Yeon and Fiona Dematraz for help analyzing data. We thank Lorenzo Scutteri for the pRD RAD9 plasmid. We thank Susan Gasser for feedback on the preliminary results. Support for ML, VG, and SJR was provided by SNSF grants CRSK-3 190526, 310030 204938, and CRSK-3 221036 awarded to SJR.

## Author contributions

ML and SJR designed the experiments. ML constructed the plasmid and the strains, and performed the experiments. ML and VG analyzed the data. ML and SJR wrote the manuscript. SJR supervised the work and acquired funding.

## Supplementary Material

### Supplementary Figures

**Supplementary Figure 1:**
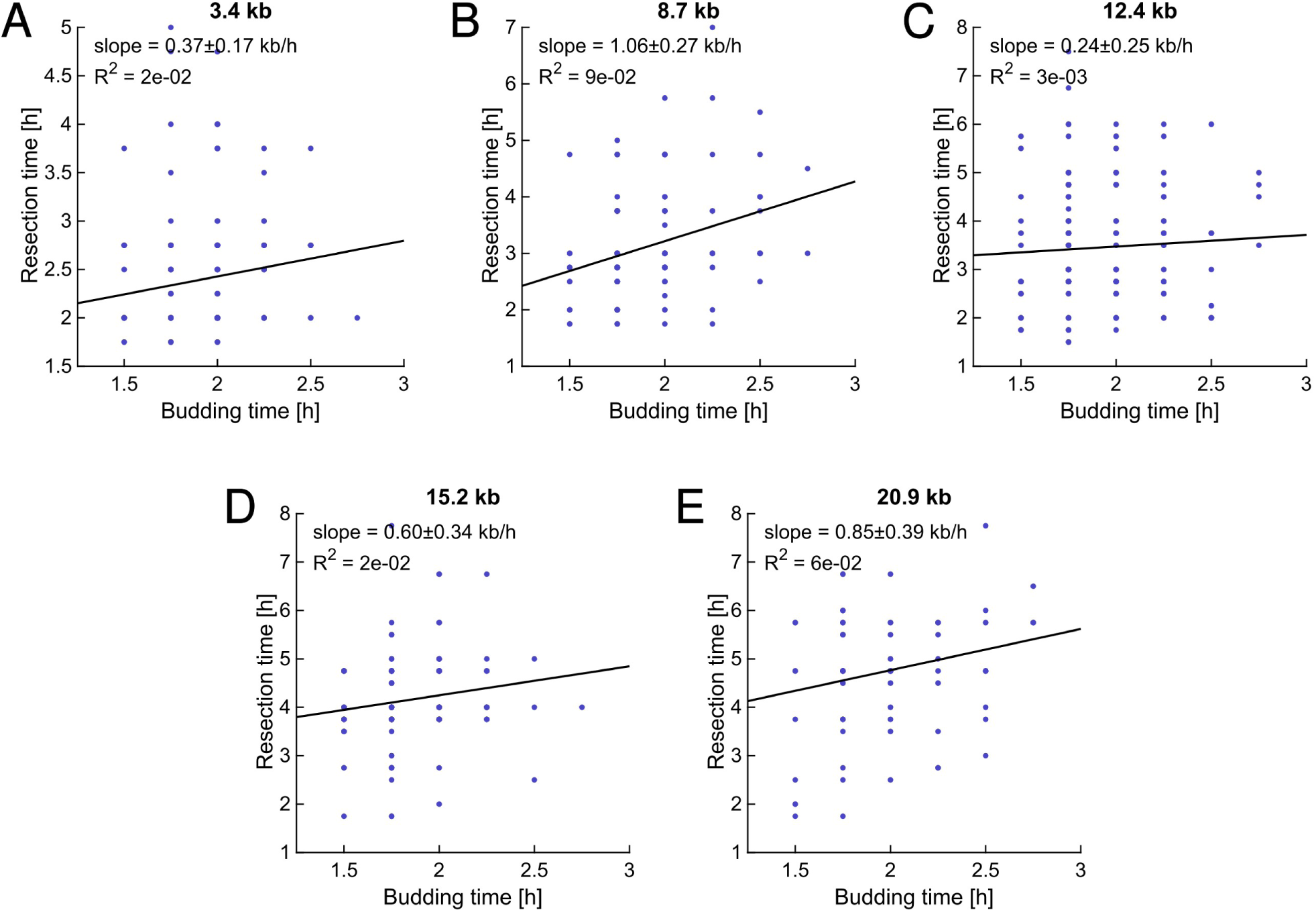
Correlation of resection times and budding times for all tripwire distances in the presence of one DSB. The oblique black lines represent linear fits.

**Supplementary Figure 2:**
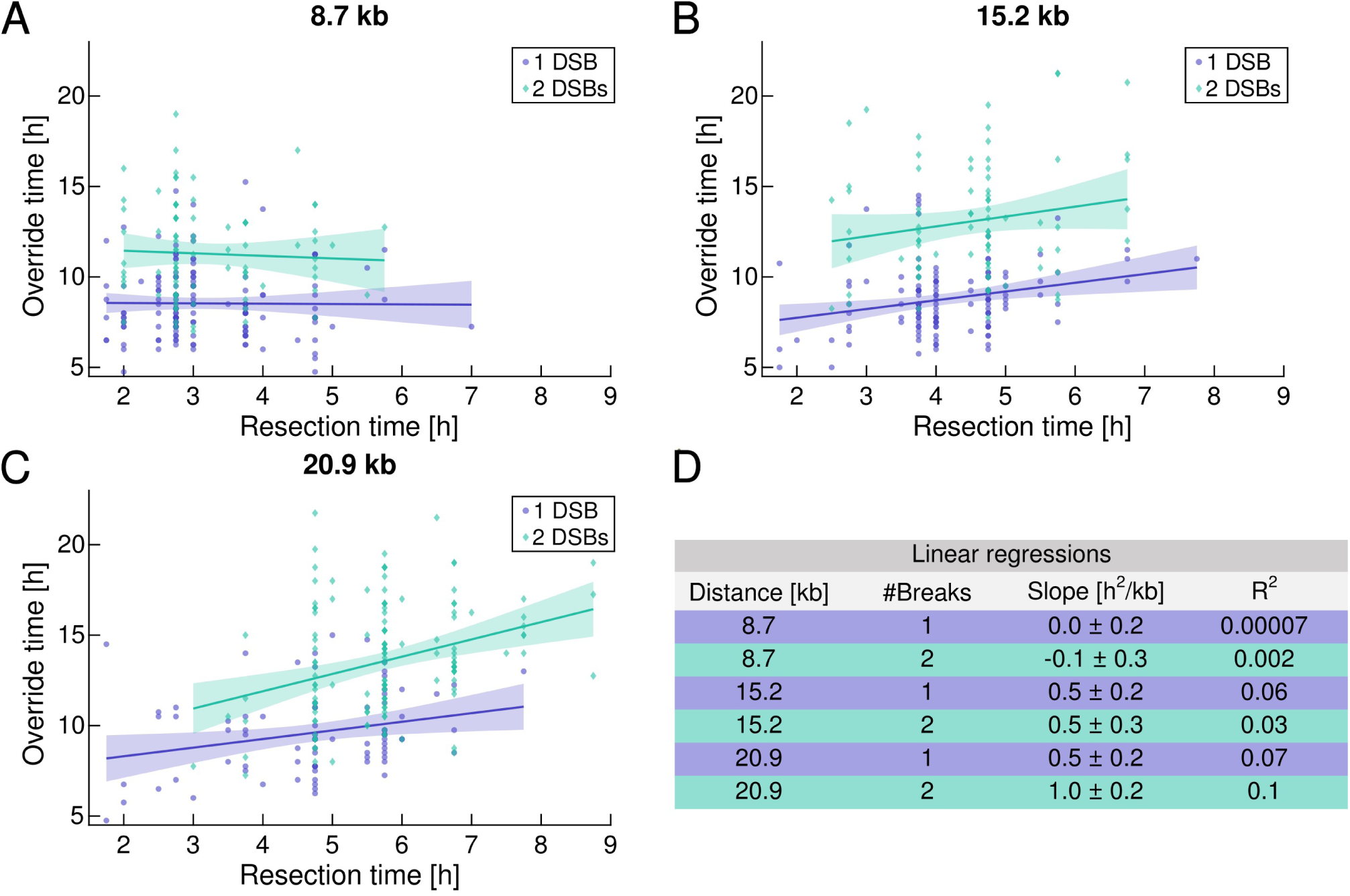
Correlations of override time and resection time. A, B, C. Override time as a function of resection time for cells with 1 or 2 DSBs with the tripwire placed at 8.7 kb (A), 15.2 kb (B) and 20.9 kb (C) from the Ho cut-site in the *MIC60* promoter. Linear regression shown with 95% confidence intervals highlighted. D. Characteristics of the linear regressions. For all panels, the number of cells as well as the override ratios are provided in Supplementary Table 1.

### Supplementary Tables

**Supplementary Table 1:**
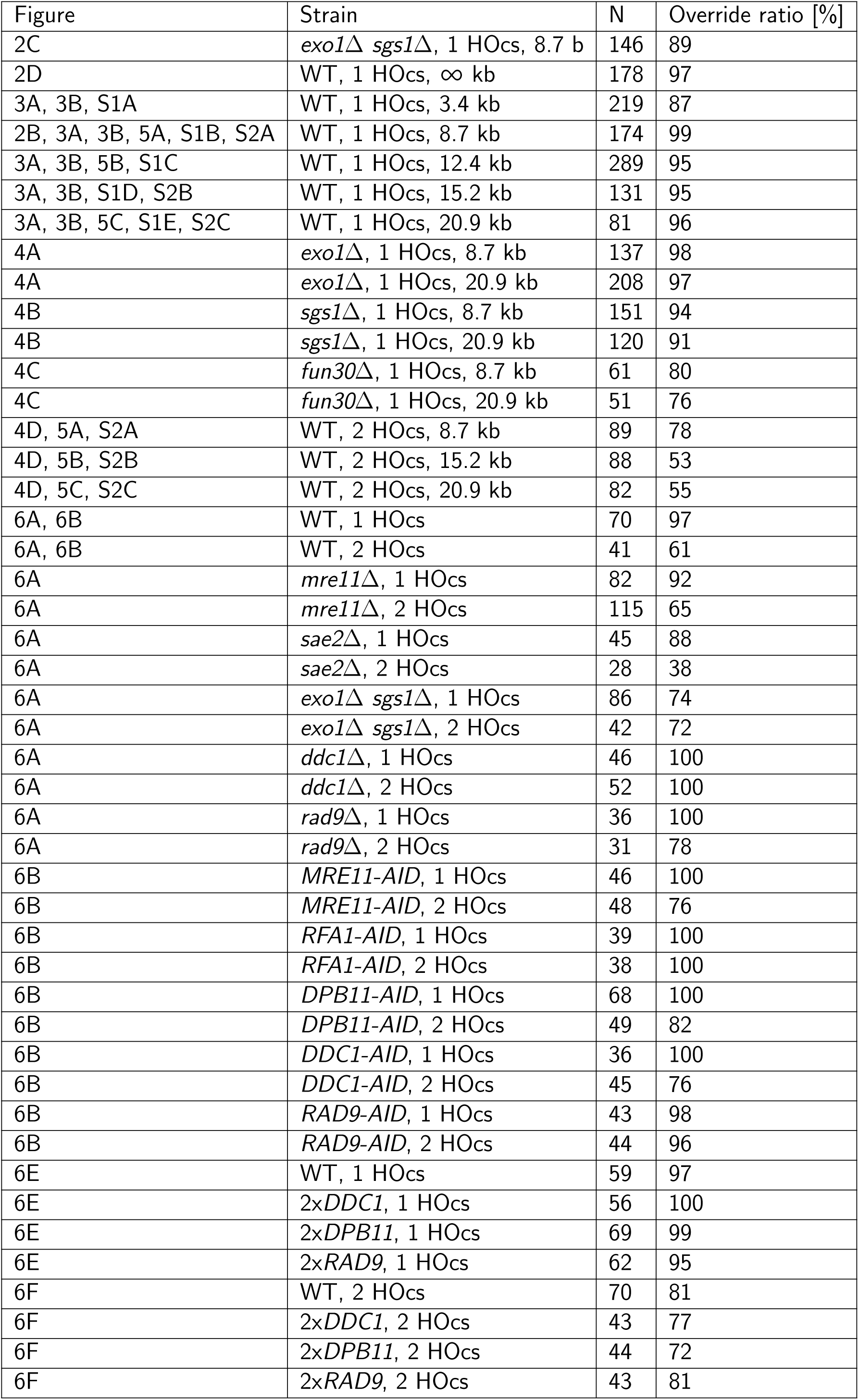
Number of cells analyzed and override ratio (percentage of cells able to override the DDC).

| Figure | Strain | N | Override ratio [%] |
| --- | --- | --- | --- |
| 2C | *exo1Δ sgs1Δ*, 1 HOcs, 8.7 kb | 146 | 89 |
| 2D | WT, 1 HOcs, ∞ kb | 178 | 97 |
| 3A, 3B, S1A | WT, 1 HOcs, 3.4 kb | 219 | 87 |
| 2B, 3A, 3B, 5A, S1B, S2A | WT, 1 HOcs, 8.7 kb | 174 | 99 |
| 3A, 3B, 5B, S1C | WT, 1 HOcs, 12.4 kb | 289 | 95 |
| 3A, 3B, S1D, S2B | WT, 1 HOcs, 15.2 kb | 131 | 95 |
| 3A, 3B, 5C, S1E, S2C | WT, 1 HOcs, 20.9 kb | 81 | 96 |
| 4A | *exo1Δ*, 1 HOcs, 8.7 kb | 137 | 98 |
| 4A | *exo1Δ*, 1 HOcs, 20.9 kb | 208 | 97 |
| 4B | *sgs1Δ*, 1 HOcs, 8.7 kb | 151 | 94 |
| 4B | *sgs1Δ*, 1 HOcs, 20.9 kb | 120 | 91 |
| 4C | *fun30Δ*, 1 HOcs, 8.7 kb | 61 | 80 |
| 4C | *fun30Δ*, 1 HOcs, 20.9 kb | 51 | 76 |
| 4D, 5A, S2A | WT, 2 HOcs, 8.7 kb | 89 | 78 |
| 4D, 5B, S2B | WT, 2 HOcs, 15.2 kb | 88 | 53 |
| 4D, 5C, S2C | WT, 2 HOcs, 20.9 kb | 82 | 55 |
| 6A, 6B | WT, 1 HOcs | 70 | 97 |
| 6A, 6B | WT, 2 HOcs | 41 | 61 |
| 6A | *mre11Δ*, 1 HOcs | 82 | 92 |
| 6A | *mre11Δ*, 2 HOcs | 115 | 65 |
| 6A | *sae2Δ*, 1 HOcs | 45 | 88 |
| 6A | *sae2Δ*, 2 HOcs | 28 | 38 |
| 6A | *exo1Δ sgs1Δ*, 1 HOcs | 86 | 74 |
| 6A | *exo1Δ sgs1Δ*, 2 HOcs | 42 | 72 |
| 6A | *ddc1Δ*, 1 HOcs | 46 | 100 |
| 6A | *ddc1Δ*, 2 HOcs | 52 | 100 |
| 6A | *rad9Δ*, 1 HOcs | 36 | 100 |
| 6A | *rad9Δ*, 2 HOcs | 31 | 78 |
| 6B | *MRE11-AID*, 1 HOcs | 46 | 100 |
| 6B | *MRE11-AID*, 2 HOcs | 48 | 76 |
| 6B | *RFA1-AID*, 1 HOcs | 39 | 100 |
| 6B | *RFA1-AID*, 2 HOcs | 38 | 100 |
| 6B | *DPB11-AID*, 1 HOcs | 68 | 100 |
| 6B | *DPB11-AID*, 2 HOcs | 49 | 82 |
| 6B | *DDC1-AID*, 1 HOcs | 36 | 100 |
| 6B | *DDC1-AID*, 2 HOcs | 45 | 76 |
| 6B | *RAD9-AID*, 1 HOcs | 43 | 98 |
| 6B | *RAD9-AID*, 2 HOcs | 44 | 96 |
| 6E | WT, 1 HOcs | 59 | 97 |
| 6E | 2x*DDC1*, 1 HOcs | 56 | 100 |
| 6E | 2x*DPB11*, 1 HOcs | 69 | 99 |
| 6E | 2x*RAD9*, 1 HOcs | 62 | 95 |
| 6F | WT, 2 HOcs | 70 | 81 |
| 6F | 2x*DDC1*, 2 HOcs | 43 | 77 |
| 6F | 2x*DPB11*, 2 HOcs | 44 | 72 |
| 6F | 2x*RAD9*, 2 HOcs | 43 | 81 |

**Supplementary Table 2:** Plasmids used in this study. All of the plasmids use *AmpR* as a bacterial selection marker.

| Plasmid name | Insert | Type of plasmid | Locus of integration | Yeast selection marker |
| --- | --- | --- | --- | --- |
| pML4 | *LIP-yEVENUS-CLN2PEST-ADH1tr*,<br>*PGK1pr-EL222-CYC1tr* | For integration | *chrXI*:466485.<br>.467107 | *natMX6* |
| pML8 | *LIP-yEVENUS-CLN2PEST-ADH1tr*,<br>*PGK1pr-EL222-CYC1tr* | For integration | *chrXI*:451520.<br>.452143 | *natMX6* |
| pML11 | *EXO1 3' UTR and EXO1 5'UTR* | For ORF deletion<br>by pop-in/pop-out | *EXO1* | *URA3*,<br>*LEU2* |
| pML12 | *SGS1 3' UTR and SGS1 5'UTR* | For ORF deletion<br>by pop-in/pop-out | *SGS1* | *URA3*,<br>*LEU2* |
| pML17 | *DPB11-yEVENUS-AID\** | For integration by<br>pop-in/pop-out | *DPB11* | *URA3*,<br>*LEU2* |
| pML20 | *MRE11 3' UTR and MRE11 5'UTR* | For ORF deletion<br>by pop-in/pop-out | *MRE11* | *URA3*,<br>*LEU2* |
| pML23 | *FUN30 3' UTR and FUN30 5'UTR* | For ORF deletion<br>by pop-in/pop-out | *FUN30* | *URA3*,<br>*LEU2* |
| pML41 | *PGK1pr-rtTA-CYC1tr* *tetOpr-*<br>*TIR1F74G-ADH1tr* | For integration in a<br>single copy | *YCR051W* | *TRP1* |
| pML48 | *MET3pr-AID\*-ymScarletI-ADH1tr* | For integration | *URA3* | *URA3* |
| pML64 | *LIP-yEVENUS-CLN2PEST-ADH1tr*,<br>*PGK1pr-EL222-CYC1tr* | For integration in a<br>single copy | *chrXI*:462286.<br>.463231 | *natMX6* |
| pML65 | *LIP-yEVENUS-CLN2PEST-ADH1tr*,<br>*PGK1pr-EL222-CYC1tr* | For integration in a<br>single copy | *chrXI*:465584.<br>.466357 | *natMX6* |
| pML66 | *LIP-yEVENUS-CLN2PEST-ADH1tr*,<br>*PGK1pr-EL222-CYC1tr* | For integration in a<br>single copy | *chrXI*:453824.<br>.454665 | *natMX6* |
| pML67 | *LIP-yEVENUS-CLN2PEST-ADH1tr*,<br>*PGK1pr-EL222-CYC1tr* | For integration in a<br>single copy | *chrXI*:453824.<br>.454665 | *natMX6* |
| pML83 | *RFA1-AID\** | For integration by<br>pop-in/pop-out | *RFA1* | *URA3*,<br>*LEU2* |
| pML84 | *MRE11-AID\** | For integration by<br>pop-in/pop-out | *MRE11* | *URA3* |
| pML85 | *SAE2 3' UTR and SAE2 5'UTR* | For ORF deletion<br>by pop-in/pop-out | *SAE2* | *URA3*,<br>*LEU2* |
| pML86 | *DDC1-AID\** | For integration by<br>pop-in/pop-out | *DDC1* | *URA3* |
| pML87 | *DDC1 3' UTR and DDC1 5'UTR* | For ORF deletion<br>by pop-in/pop-out | *DDC1* | *URA3*,<br>*LEU2* |
| pML88 | *RAD9-AID\** | For integration by<br>pop-in/pop-out | *RAD9* | *URA3*,<br>*LEU2* |
| pML89 | *RAD9 3' UTR and RAD9 5'UTR* | For ORF deletion<br>by pop-in/pop-out | *RAD9* | *URA3*,<br>*LEU2* |
| pML98 | *DDC1pr* (255 bp) - *DDC1* - *DDCtr*<br>(287 bp) | For integration in a<br>single copy | *YCR051W* | *URA3* |
| pML99 | *DPB11pr* (874 bp) - *DPB11* -<br>*DPB11tr* (180 bp) | For integration in a<br>single copy | *YCR051W* | *URA3* |
| pRD50<br>_RAD9 | *RAD9pr* (301 bp) - *RAD9* - *RAD9tr*<br>(300 bp) | For integration in a<br>single copy | *YCR051W* | *URA3* |

**Supplementary Table 3:**
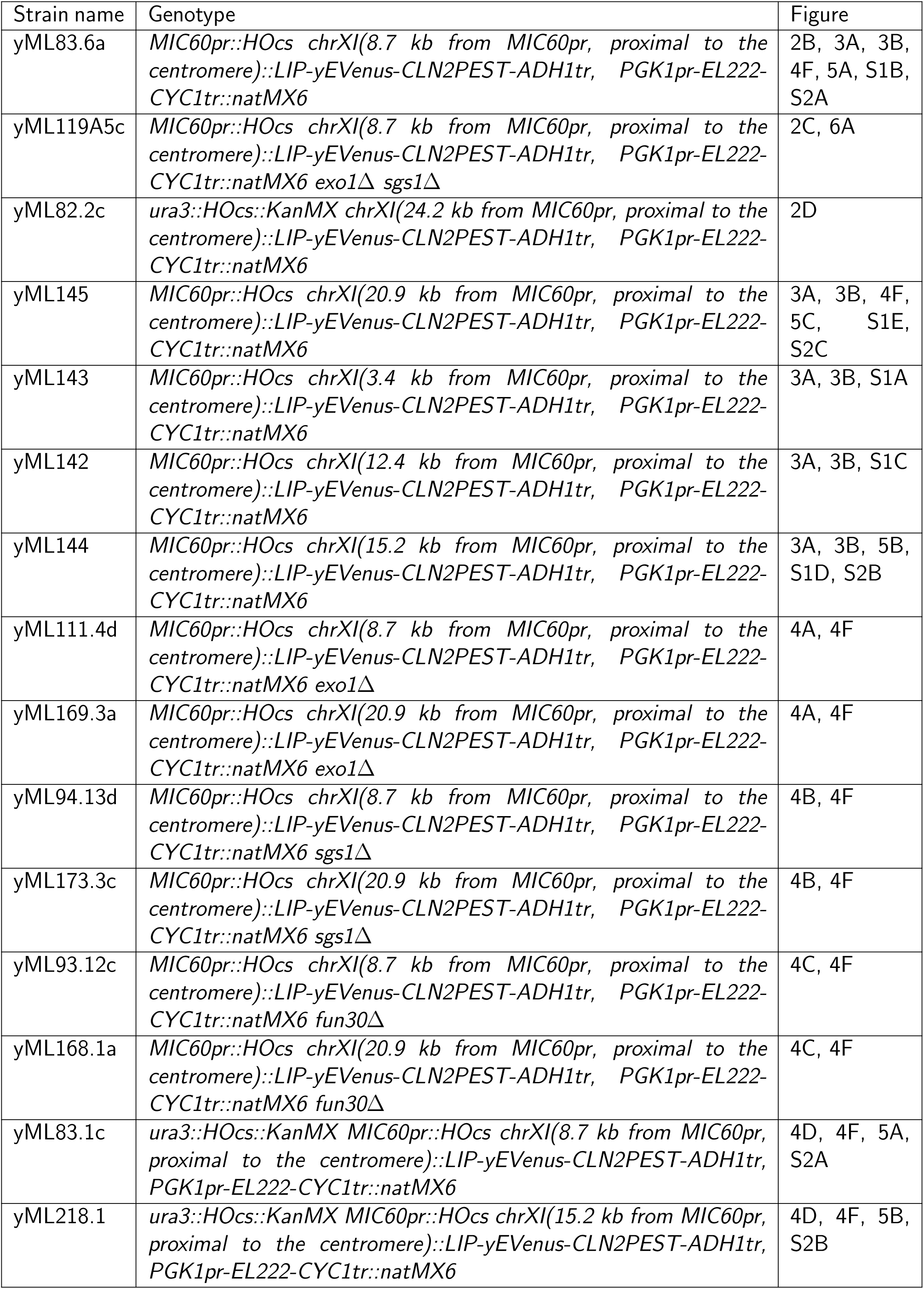
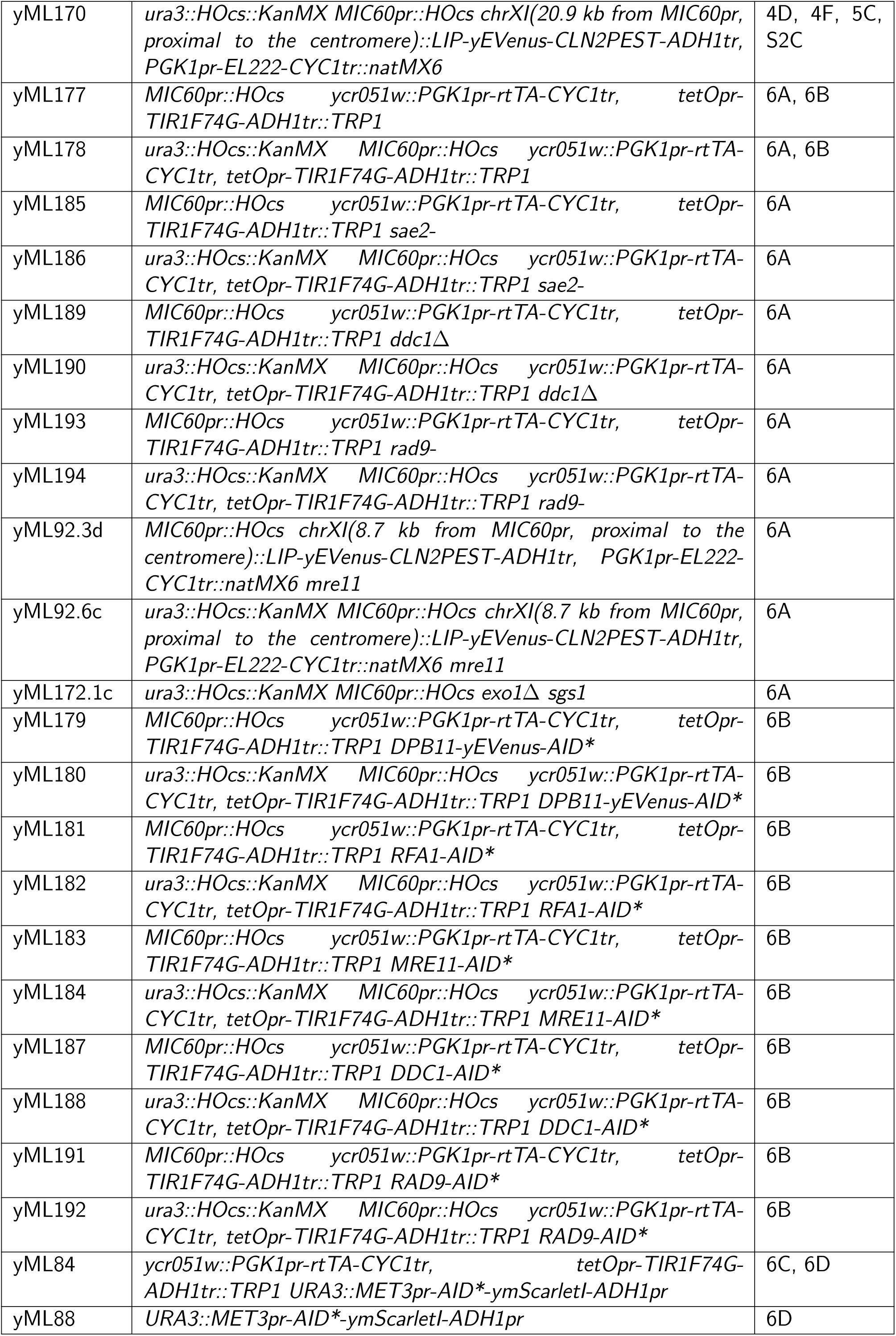
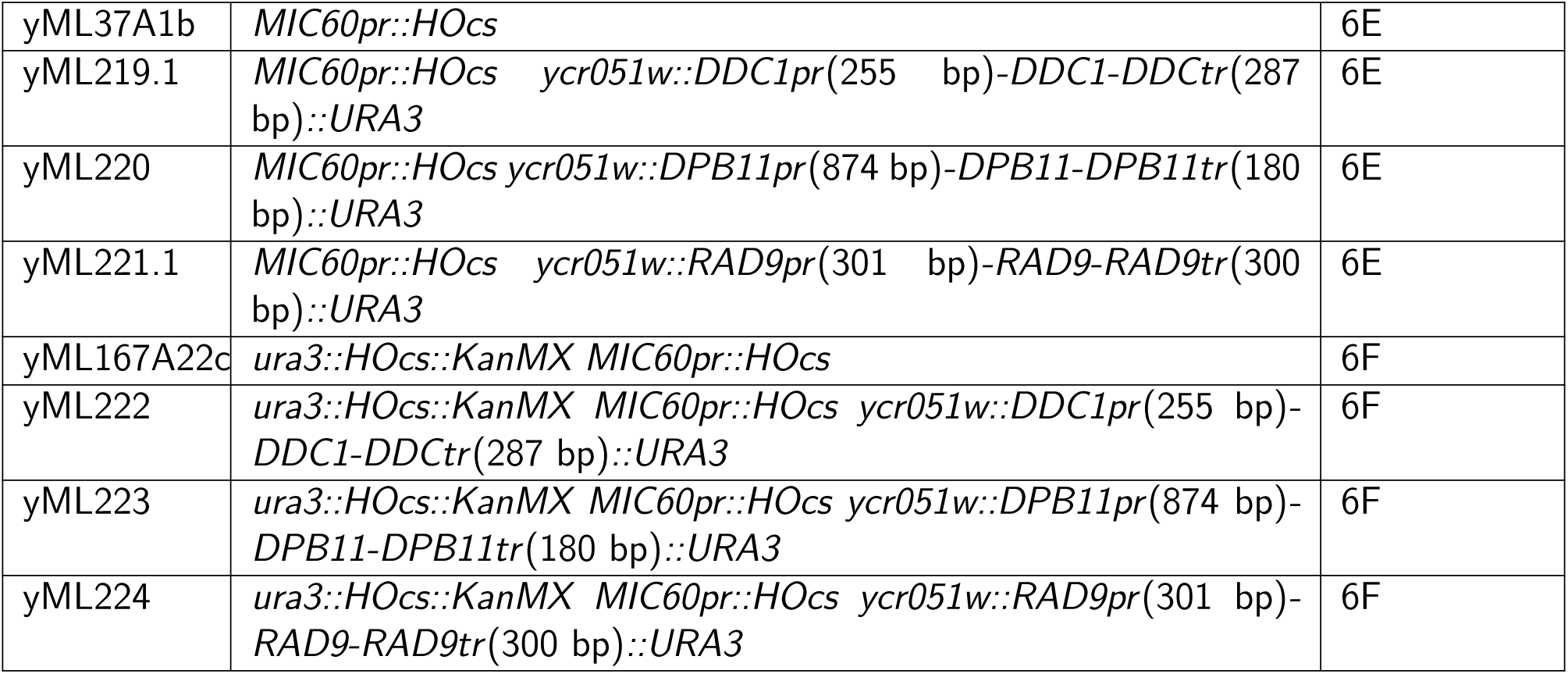
Strains used for this study. Except when indicated by an asterix (for yML84 and yML88), all strains have the common genotype *MATα-syn cln1*Δ *cln2::MET3pr-CLN2 cln3::GAL1pr-HO HTB2-mCherry::HIS5 RAD5-corrected*, which we omitted for clarity.

| Strain name | Genotype | Figure |
| --- | --- | --- |
| yML83.6a | *MIC60pr::HOcs chrXI(8.7 kb from MIC60pr, proximal to the centromere)::LIP-yEVenus-CLN2PEST-ADH1tr, PGK1pr-EL222-CYC1tr::natMX6* | 2B, 3A, 3B, 4F, 5A, S1B, S2A |
| yML119A5c | *MIC60pr::HOcs chrXI(8.7 kb from MIC60pr, proximal to the centromere)::LIP-yEVenus-CLN2PEST-ADH1tr, PGK1pr-EL222-CYC1tr::natMX6 exo1Δ sgs1Δ* | 2C, 6A |
| yML82.2c | *ura3::HOcs::KanMX chrXI(24.2 kb from MIC60pr, proximal to the centromere)::LIP-yEVenus-CLN2PEST-ADH1tr, PGK1pr-EL222-CYC1tr::natMX6* | 2D |
| yML145 | *MIC60pr::HOcs chrXI(20.9 kb from MIC60pr, proximal to the centromere)::LIP-yEVenus-CLN2PEST-ADH1tr, PGK1pr-EL222-CYC1tr::natMX6* | 3A, 3B, 4F, 5C, S1E, S2C |
| yML143 | *MIC60pr::HOcs chrXI(3.4 kb from MIC60pr, proximal to the centromere)::LIP-yEVenus-CLN2PEST-ADH1tr, PGK1pr-EL222-CYC1tr::natMX6* | 3A, 3B, S1A |
| yML142 | *MIC60pr::HOcs chrXI(12.4 kb from MIC60pr, proximal to the centromere)::LIP-yEVenus-CLN2PEST-ADH1tr, PGK1pr-EL222-CYC1tr::natMX6* | 3A, 3B, S1C |
| yML144 | *MIC60pr::HOcs chrXI(15.2 kb from MIC60pr, proximal to the centromere)::LIP-yEVenus-CLN2PEST-ADH1tr, PGK1pr-EL222-CYC1tr::natMX6* | 3A, 3B, 5B, S1D, S2B |
| yML111.4d | *MIC60pr::HOcs chrXI(8.7 kb from MIC60pr, proximal to the centromere)::LIP-yEVenus-CLN2PEST-ADH1tr, PGK1pr-EL222-CYC1tr::natMX6 exo1Δ* | 4A, 4F |
| yML169.3a | *MIC60pr::HOcs chrXI(20.9 kb from MIC60pr, proximal to the centromere)::LIP-yEVenus-CLN2PEST-ADH1tr, PGK1pr-EL222-CYC1tr::natMX6 exo1Δ* | 4A, 4F |
| yML94.13d | *MIC60pr::HOcs chrXI(8.7 kb from MIC60pr, proximal to the centromere)::LIP-yEVenus-CLN2PEST-ADH1tr, PGK1pr-EL222-CYC1tr::natMX6 sgs1Δ* | 4B, 4F |
| yML173.3c | *MIC60pr::HOcs chrXI(20.9 kb from MIC60pr, proximal to the centromere)::LIP-yEVenus-CLN2PEST-ADH1tr, PGK1pr-EL222-CYC1tr::natMX6 sgs1Δ* | 4B, 4F |
| yML93.12c | *MIC60pr::HOcs chrXI(8.7 kb from MIC60pr, proximal to the centromere)::LIP-yEVenus-CLN2PEST-ADH1tr, PGK1pr-EL222-CYC1tr::natMX6 fun30Δ* | 4C, 4F |
| yML168.1a | *MIC60pr::HOcs chrXI(20.9 kb from MIC60pr, proximal to the centromere)::LIP-yEVenus-CLN2PEST-ADH1tr, PGK1pr-EL222-CYC1tr::natMX6 fun30Δ* | 4C, 4F |
| yML83.1c | *ura3::HOcs::KanMX MIC60pr::HOcs chrXI(8.7 kb from MIC60pr, proximal to the centromere)::LIP-yEVenus-CLN2PEST-ADH1tr, PGK1pr-EL222-CYC1tr::natMX6* | 4D, 4F, 5A, S2A |
| yML218.1 | *ura3::HOcs::KanMX MIC60pr::HOcs chrXI(15.2 kb from MIC60pr, proximal to the centromere)::LIP-yEVenus-CLN2PEST-ADH1tr, PGK1pr-EL222-CYC1tr::natMX6* | 4D, 4F, 5B, S2B |
| yML170 | *ura3::HOcs::KanMX MIC60pr::HOcs chrXI(20.9 kb from MIC60pr, proximal to the centromere)::LIP-yEVENUS-CLN2PEST-ADH1tr, PGK1pr-EL222-CYC1tr::natMX6* | 4D, 4F, 5C, S2C |
| yML177 | *MIC60pr::HOcs ycr051w::PGK1pr-rtTA-CYC1tr, tetOpr-TIR1F74G-ADH1tr::TRP1* | 6A, 6B |
| yML178 | *ura3::HOcs::KanMX MIC60pr::HOcs ycr051w::PGK1pr-rtTA-CYC1tr, tetOpr-TIR1F74G-ADH1tr::TRP1* | 6A, 6B |
| yML185 | *MIC60pr::HOcs ycr051w::PGK1pr-rtTA-CYC1tr, tetOpr-TIR1F74G-ADH1tr::TRP1 sae2-* | 6A |
| yML186 | *ura3::HOcs::KanMX MIC60pr::HOcs ycr051w::PGK1pr-rtTA-CYC1tr, tetOpr-TIR1F74G-ADH1tr::TRP1 sae2-* | 6A |
| yML189 | *MIC60pr::HOcs ycr051w::PGK1pr-rtTA-CYC1tr, tetOpr-TIR1F74G-ADH1tr::TRP1 ddc1Δ* | 6A |
| yML190 | *ura3::HOcs::KanMX MIC60pr::HOcs ycr051w::PGK1pr-rtTA-CYC1tr, tetOpr-TIR1F74G-ADH1tr::TRP1 ddc1Δ* | 6A |
| yML193 | *MIC60pr::HOcs ycr051w::PGK1pr-rtTA-CYC1tr, tetOpr-TIR1F74G-ADH1tr::TRP1 rad9-* | 6A |
| yML194 | *ura3::HOcs::KanMX MIC60pr::HOcs ycr051w::PGK1pr-rtTA-CYC1tr, tetOpr-TIR1F74G-ADH1tr::TRP1 rad9-* | 6A |
| yML92.3d | *MIC60pr::HOcs chrXI(8.7 kb from MIC60pr, proximal to the centromere)::LIP-yEVENUS-CLN2PEST-ADH1tr, PGK1pr-EL222-CYC1tr::natMX6 mre11* | 6A |
| yML92.6c | *ura3::HOcs::KanMX MIC60pr::HOcs chrXI(8.7 kb from MIC60pr, proximal to the centromere)::LIP-yEVENUS-CLN2PEST-ADH1tr, PGK1pr-EL222-CYC1tr::natMX6 mre11* | 6A |
| yML172.1c | *ura3::HOcs::KanMX MIC60pr::HOcs exo1Δ sgs1* | 6A |
| yML179 | *MIC60pr::HOcs ycr051w::PGK1pr-rtTA-CYC1tr, tetOpr-TIR1F74G-ADH1tr::TRP1 DPB11-yEVENUS-AID\** | 6B |
| yML180 | *ura3::HOcs::KanMX MIC60pr::HOcs ycr051w::PGK1pr-rtTA-CYC1tr, tetOpr-TIR1F74G-ADH1tr::TRP1 DPB11-yEVENUS-AID\** | 6B |
| yML181 | *MIC60pr::HOcs ycr051w::PGK1pr-rtTA-CYC1tr, tetOpr-TIR1F74G-ADH1tr::TRP1 RFA1-AID\** | 6B |
| yML182 | *ura3::HOcs::KanMX MIC60pr::HOcs ycr051w::PGK1pr-rtTA-CYC1tr, tetOpr-TIR1F74G-ADH1tr::TRP1 RFA1-AID\** | 6B |
| yML183 | *MIC60pr::HOcs ycr051w::PGK1pr-rtTA-CYC1tr, tetOpr-TIR1F74G-ADH1tr::TRP1 MRE11-AID\** | 6B |
| yML184 | *ura3::HOcs::KanMX MIC60pr::HOcs ycr051w::PGK1pr-rtTA-CYC1tr, tetOpr-TIR1F74G-ADH1tr::TRP1 MRE11-AID\** | 6B |
| yML187 | *MIC60pr::HOcs ycr051w::PGK1pr-rtTA-CYC1tr, tetOpr-TIR1F74G-ADH1tr::TRP1 DDC1-AID\** | 6B |
| yML188 | *ura3::HOcs::KanMX MIC60pr::HOcs ycr051w::PGK1pr-rtTA-CYC1tr, tetOpr-TIR1F74G-ADH1tr::TRP1 DDC1-AID\** | 6B |
| yML191 | *MIC60pr::HOcs ycr051w::PGK1pr-rtTA-CYC1tr, tetOpr-TIR1F74G-ADH1tr::TRP1 RAD9-AID\** | 6B |
| yML192 | *ura3::HOcs::KanMX MIC60pr::HOcs ycr051w::PGK1pr-rtTA-CYC1tr, tetOpr-TIR1F74G-ADH1tr::TRP1 RAD9-AID\** | 6B |
| yML84 | *ycr051w::PGK1pr-rtTA-CYC1tr, tetOpr-TIR1F74G-ADH1tr::TRP1 URA3::MET3pr-AID\*-ymScarletI-ADH1pr* | 6C, 6D |
| yML88 | *URA3::MET3pr-AID\*-ymScarletI-ADH1pr* | 6D |
| yML37A1b | *MIC60pr::HOcs* | 6E |
| yML219.1 | *MIC60pr::HOcs ycr051w::DDC1pr(255 bp)-DDC1-DDCtr(287 bp)::URA3* | 6E |
| yML220 | *MIC60pr::HOcs ycr051w::DPB11pr(874 bp)-DPB11-DPB11tr(180 bp)::URA3* | 6E |
| yML221.1 | *MIC60pr::HOcs ycr051w::RAD9pr(301 bp)-RAD9-RAD9tr(300 bp)::URA3* | 6E |
| yML167A22c | *ura3::HOcs::KanMX MIC60pr::HOcs* | 6F |
| yML222 | *ura3::HOcs::KanMX MIC60pr::HOcs ycr051w::DDC1pr(255 bp)-DDC1-DDCtr(287 bp)::URA3* | 6F |
| yML223 | *ura3::HOcs::KanMX MIC60pr::HOcs ycr051w::DPB11pr(874 bp)-DPB11-DPB11tr(180 bp)::URA3* | 6F |
| yML224 | *ura3::HOcs::KanMX MIC60pr::HOcs ycr051w::RAD9pr(301 bp)-RAD9-RAD9tr(300 bp)::URA3* | 6F |

## References

1. Negrini, S., Gorgoulis, V. G. & Halazonetis, T. D. Genomic instability – an evolving hallmark of cancer. Nat. Rev. Mol. Cell Biol. 11, 220–228 (2010).

2. Malkova, A. & Haber, J. E. Mutations arising during repair of chromosome breaks. Annu. Rev. Genet. 46, 455–473 (2012).

3. Waterman, D. P., Haber, J. E. & Smolka, M. B. Checkpoint responses to DNA double-strand breaks. Annu. Rev. Biochem. 89, 103–133 (2020).

4. Symington, L. S. & Gautier, J. Double-strand break end resection and repair pathway choice. Annu. Rev. Genet. 45, 247–271 (2011).

5. Oh, J., Al-Zain, A., Cannavo, E., Cejka, P. & Symington, L. S. Xrs2 dependent and independent functions of the Mre11-Rad50 complex. Mol. Cell 64, 405–415 (2016).

6. Andres, S. N. & Williams, R. S. CtIP/Ctp1/Sae2, molecular form fit for function. DNA Repair 56, 109–117 (2017). Cutting-edge Perspectives in Genomic Maintenance IV.

7. Mimitou, E. P. & Symington, L. S. Sae2, Exo1 and Sgs1 collaborate in DNA double-strand break processing. Nature 455, 770–774 (2008).

8. Shim, E. Y., Chung, W., Nicolette, M. L., Zhang, Y., Davis, M., Zhu, Z., Paull, T. T., Ira, G. & Lee, S. E. *Saccharomyces cerevisiae* Mre11/Rad50/Xrs2 and Ku proteins regulate association of Exo1 and Dna2 with DNA breaks. EMBO J. 29, 3370–3380 (2010).

9. Mimitou, E. P. & Symington, L. S. Ku prevents Exo1 and Sgs1-dependent resection of DNA ends in the absence of a functional MRX complex or Sae2. EMBO J. 29, 3358–3369 (2010).

10. Zhu, Z., Chung, W.-H., Shim, E. Y., Lee, S. E. & Ira, G. Sgs1 helicase and two nucleases Dna2 and Exo1 resect DNA double-strand break ends. Cell 134, 981–994 (2008).

11. Zou, L. & Elledge, S. J. Sensing DNA damage through ATRIP recognition of RPA-ssDNA complexes. Science 300, 1542–1548 (2003).

12. Sun, Z., Hsiao, J., Fay, D. S. & Stern, D. F. Rad53 FHA domain associated with phosphorylated Rad9 in the DNA damage checkpoint. Science 281, 272–274 (1998).

13. Sweeney, F. D., Yang, F., Chi, A., Shabanowitz, J., Hunt, D. F. & Durocher, D. *Saccharomyces cerevisiae* Rad9 acts as a Mec1 adaptor to allow Rad53 activation. Curr. Biol. 15, 1364–1375 (2005).

14. Jaehnig, E., Kuo, D., Hombauer, H., Ideker, T. & Kolodner, R. Checkpoint kinases regulate a global network of transcription factors in response to DNA damage. Cell Rep. 4, 174–188 (2013).

15. Sandell, L. L. & Zakian, V. A. Loss of a yeast telomere: Arrest, recovery, and chromosome loss. Cell 75, 729–739 (1993).

16. Toczyski, D. P., Galgoczy, D. J. & Hartwell, L. H. CDC5 and CKII control adaptation to the yeast DNA damage checkpoint. Cell 90, 1097–1106 (1997).

17. Syljuåsen, R. G., Jensen, S., Bartek, J. & Lukas, J. Adaptation to the ionizing radiation–induced G2 checkpoint occurs in human cells and depends on checkpoint kinase 1 and polo-like kinase 1 kinases. Cancer Res. 66, 10253–10257 (2006).

18. Syljuåsen, R. G. Checkpoint adaptation in human cells. Oncogene 26, 5833–5839 (2007).

19. Swift, L. & Golsteyn, R. Chapter 22 -the relationship between checkpoint adaptation and mitotic catastrophe in genomic changes in cancer cells. In Genome Stability (eds. Kovalchuk, I. & Kovalchuk, O.), 373–389 (Academic Press, 2016).

20. Sadeghi, A., Dervey, R., Gligorovski, V., Labagnara, M. & Rahi, S. J. The optimal strategy balancing risk and speed predicts DNA damage checkpoint override times. Nature Physics 18, 832–839 (2022).

21. Lee, S. E., Moore, J., Holmes, A., Umezu, K., Kolodner, R. D. & Haber, J. E. *Saccharomyces* Ku70, Mre11/Rad50, and RPA proteins regulate adaptation to G2/M arrest after DNA damage. Cell 94, 399–409 (1998).

22. Pellicioli, A., Lee, S. E., Lucca, C., Foiani, M. & Haber, J. E. Regulation of *Saccharomyces* Rad53 checkpoint kinase during adaptation from DNA damage–induced G2/M arrest. Mol. Cell 7, 293–300 (2001).

23. Mantiero, D., Clerici, M., Lucchini, G. & Longhese, M. P. Dual role for *Saccharomyces cerevisiae* Tel1 in the checkpoint response to double-strand breaks. EMBO Rep. 8, 380–387 (2007).

24. Jazayeri, A., Balestrini, A., Garner, E., Haber, J. E. & Costanzo, V. Mre11–Rad50–Nbs1-dependent processing of DNA breaks generates oligonucleotides that stimulate ATM activity. EMBO J. 27, 1953–1962 (2008).

25. Eapen, V. V., Sugawara, N., Tsabar, M., Wu, W.-H. & Haber, J. E. The *Saccharomyces cerevisiae* chromatin remodeler Fun30 regulates DNA end resection and checkpoint deactivation. Mol. Cell. Biol. 32, 4727–4740 (2012).

26. White, C. & Haber, J. Intermediates of recombination during mating type switching in *Saccharomyces cerevisiae*. EMBO J. 9, 663–673 (1990).

27. Cao, L., Alani, E. & Kleckner, N. A pathway for generation and processing of double-strand breaks during meiotic recombination in *S. cerevisiae*. Cell 61, 1089–1101 (1990).

28. Zierhut, C. & Diffley, J. F. X. Break dosage, cell cycle stage and DNA replication influence DNA double strand break response. EMBO J. 27, 1875–1885 (2008).

29. Bazzano, D., Lomonaco, S. & Wilson, T. E. Mapping yeast mitotic 5’ resection at base resolution reveals the sequence and positional dependence of nucleases *in vivo*. Nucleic Acids Res. 49, 12607–12621 (2021).

30. Whelan, D. R. & Rothenberg, E. Super-resolution mapping of cellular double-strand break resection complexes during homologous recombination. Proc. Natl. Acad. Sci. U. S. A. 118, e2021963118 (2021).

31. Leland, B. A., Chen, A. C., Zhao, A. Y., Wharton, R. C. & King, M. C. Rev7 and 53BP1/Crb2 prevent RecQ helicase-dependent hyper-resection of DNA double-strand breaks. eLife 7, e33402 (2018).

32. Manfrini, N., Clerici, M., Wery, M., Colombo, C. V., Descrimes, M., Morillon, A., d’Adda di Fagagna, F. & Longhese, M. P. Resection is responsible for loss of transcription around a double-strand break in *Saccharomyces cerevisiae*. eLife 4, e08942 (2015).

33. Rivera-Cancel, G., Motta-Mena, L. B. & Gardner, K. H. Identification of natural and artificial DNA substrates for light-activated LOV–HTH transcription factor EL222. Biochemistry 51, 10024–10034 (2012). PMID: 23205774.

34. Gligorovski, V., Sadeghi, A. & Rahi, S. J. Multidimensional characterization of inducible promoters and a highly light-sensitive LOV-transcription factor. Nature Communications 14, 3810 (2023).

35. Peng, B., Williams, T. C., Henry, M., Nielsen, L. K. & Vickers, C. E. Controlling heterologous gene expression in yeast cell factories on different carbon substrates and across the diauxic shift: a comparison of yeast promoter activities. Microb. Cell Fact. 14, 91 (2015).

36. Oesterle, R. & Rahi, S. J. Why budding yeast overrides the dna damage checkpoint. bioRxiv 2025.07.30.667764 (2025).

37. Amon, A., Irniger, S. & Nasmyth, K. Closing the cell cycle circle in yeast: G2 cyclin proteolysis initiated at mitosis persists until the activation of G1 cyclins in the next cycle. Cell 77, 1037–1050 (1994).

38. Kaboli, S., Yamakawa, T., Sunada, K., Takagaki, T., Sasano, Y., Sugiyama, M., Kaneko, Y. & Harashima, S. Genome-wide mapping of unexplored essential regions in the *Saccharomyces cerevisiae* genome: evidence for hidden synthetic lethal combinations in a genetic interaction network. Nucleic Acids Res. 42, 9838–9853 (2014).

39. Clemente-Blanco, A., Mayán-Santos, M., Schneider, D. A., Machín, F., Jarmuz, A., Tschochner, H. & Aragón, L. Cdc14 inhibits transcription by RNA polymerase I during anaphase. Nature 458, 219–222 (2009).

40 . Memişoǧlu, G., Bohn, S., Krogan, N. J., Haber, J. E. & Ruthenburg, A. J. The mediator kinase module regulates cell cycle re-entry and transcriptional responses following DNA damage. bioRxiv 2023.02.26.530133 (2023).

41. Di Talia, S., Skotheim, J. M., Bean, J. M., Siggia, E. D. & Cross, F. R. The effects of molecular noise and size control on variability in the budding yeast cell cycle. Nature 448, 947–951 (2007).

42. Fishman-Lobell, J., Rudin, N. & Haber, J. E. Two alternative pathways of double-strand break repair that are kinetically separable and independently modulated. Mol. Cell. Biol. 12, 1292– 1303 (1992).

43. Cassani, C., Gobbini, E., Vertemara, J., Wang, W., Marsella, A., Sung, P., Tisi, R., Zampella, G. & Longhese, M. P. Structurally distinct Mre11 domains mediate MRX functions in resection, end-tethering and DNA damage resistance. Nucleic Acids Res. 46, 2990–3008 (2018).

44. Mordes, D. A., Nam, E. A. & Cortez, D. Dpb11 activates the Mec1–Ddc2 complex. Proc. Natl. Acad. Sci. U.S.A. 105, 18730–18734 (2008).

45. Navadgi-Patil, V. M. & Burgers, P. M. Yeast DNA replication protein Dpb11 activates the Mec1/ATR checkpoint kinase. J. Biol. Chem. 283, 35853–35859 (2008).

46. Granata, M., Lazzaro, F., Novarina, D., Panigada, D., Puddu, F., Abreu, C. M., Kumar, R., Grenon, M., Lowndes, N. F., Plevani, P. & Muzi-Falconi, M. Dynamics of Rad9 chromatin binding and checkpoint function are mediated by its dimerization and are cell cycle-regulated by CDK1 activity. PLoS Genet. 6, e1001047 (2010).

47. Pfander, B. & Diffley, J. F. X. Dpb11 coordinates Mec1 kinase activation with cell cycle-regulated Rad9 recruitment. EMBO J. 30, 4897–4907 (2011).

48. Puddu, F., Granata, M., Nola, L. D., Balestrini, A., Piergiovanni, G., Lazzaro, F., Giannattasio, M., Plevani, P. & Muzi-Falconi, M. Phosphorylation of the budding yeast 9-1-1 complex is required for Dpb11 function in the full activation of the UV-induced DNA damage checkpoint. Mol. Cell. Biol. 28, 4782–4793 (2008). PMID: 18541674.

49. Usui, T., Foster, S. S. & Petrini, J. H. Maintenance of the DNA-damage checkpoint requires DNA-damage-induced mediator protein oligomerization. Mol. Cell 33, 147–159 (2009).

50. Schwartz, M. F., Duong, J. K., Sun, Z., Morrow, J. S., Pradhan, D. & Stern, D. F. Rad9 phosphorylation sites couple Rad53 to the *Saccharomyces cerevisiae* DNA damage checkpoint. Mol. Cell 9, 1055–1065 (2002).

51. Morawska, M. & Ulrich, H. D. An expanded tool kit for the auxin-inducible degron system in budding yeast. Yeast 30, 341–351 (2013).

52. Lu, Z., Peng, B., Ebert, B. E., Dumsday, G. & Vickers, C. E. Auxin-mediated protein depletion for metabolic engineering in terpene-producing yeast. Nat. Commun. 12, 1051 (2021).

53. Yesbolatova, A., Saito, Y., Kitamoto, N., Makino-Itou, H., Ajima, R., Nakano, R., Nakaoka, H., Fukui, K., Gamo, K., Tominari, Y., Takeuchi, H., Saga, Y., Hayashi, K.-i. & Kanemaki, M. T. The auxin-inducible degron 2 technology provides sharp degradation control in yeast, mammalian cells, and mice. Nat. Commun. 11, 5701 (2020).

54. Papagiannakis, A., de Jonge, J. J., Zhang, Z. & Heinemann, M. Quantitative characterization of the auxin-inducible degron: a guide for dynamic protein depletion in single yeast cells. Sci. Rep. 7, 4704 (2017).

55. Majka, J., Binz, S. K., Wold, M. S. & Burgers, P. M. J. Replication Protein A directs loading of the DNA damage checkpoint clamp to 5’-DNA junctions. J. Biol. Chem. 281, 27855–27861 (2006).

56. Ellison, V. & Stillman, B. Biochemical characterization of DNA damage checkpoint complexes: clamp loader and clamp complexes with specificity for 5’ recessed DNA. PLoS Biol. 1, E33 (2003).

57. Lisby, M., Barlow, J. H., Burgess, R. C. & Rothstein, R. Choreography of the DNA damage response: Spatiotemporal relationships among checkpoint and repair proteins. Cell 118, 699– 713 (2004).

58. Peritore, M., Reusswig, K.-U., Bantele, S. C. S., Straub, T. & Pfander, B. Strand-specific ChIP-seq at DNA breaks distinguishes ssDNA versus dsDNA binding and refutes single-stranded nucleosomes. Mol. Cell 81, 1841–1853.e4 (2021).

59. Yu, T.-Y., Kimble, M. T. & Symington, L. S. Sae2 antagonizes Rad9 accumulation at DNA double-strand breaks to attenuate checkpoint signaling and facilitate end resection. Proc. Natl. Acad. Sci. U.S.A. 115, E11961–E11969 (2018).

60. di Cicco, G., Bantele, S. C. S., Reusswig, K.-U. & Pfander, B. A cell cycle-independent mode of the Rad9-Dpb11 interaction is induced by DNA damage. Sci. Rep. 7, 11650 (2017).

61. Gordon, A., Colman-Lerner, A., Chin, T. E., Benjamin, K. R., Yu, R. C. & Brent, R. Single-cell quantification of molecules and rates using open-source microscope-based cytometry. Nat. Methods 4, 175–181 (2007).

62. Rahi, S., Pecani, K., Ondracka, A., Oikonomou, C. & Cross, F. The CDK-APC/C oscillator predominantly entrains periodic cell-cycle transcription. Cell 165, 475–487 (2016).

63. Rahi, S. J., Larsch, J., Pecani, K., Katsov, A. Y., Mansouri, N., Tsaneva-Atanasova, K., Sontag, E. D. & Cross, F. R. Oscillatory stimuli differentiate adapting circuit topologies. Nature Methods 14, 1010–1016 (2017).

64. Gietz, R. D. & Schiestl, R. H. High-efficiency yeast transformation using the LiAc/SS carrier DNA/PEG method. Nat. Protoc. 2, 31–34 (2007).

65. Stojković, L., Gligorovski, V., Geramimanesh, M., Labagnara, M. & Rahi, S. J. Automated plasmid design for marker-free genome editing in budding yeast. G3 (Bethesda) 15, jkae297 (2024).

66. Elserafy, M. & El-Khamisy, S. F. Choose your yeast strain carefully: the RAD5 gene matters. Nat. Rev. Mol. Cell Biol. 19, 343–344 (2018).

67. Gligorovski, V., Labagnara, M., Scutteri, L., Blackholm, M., Möglich, A., Mansouri, N. & Rahi, S. J. Light-directed evolution of dynamic, multi-state, and computational protein functionalities. bioRxiv 2024.02.28.582517 (2024).

68. Dietler, N., Minder, M., Gligorovski, V., Economou, A. M., Joly, D. A. H. L., Sadeghi, A., Chan, C. H. M., Koziński, M., Weigert, M., Bitbol, A.-F. & Rahi, S. J. A convolutional neural network segments yeast microscopy images with high accuracy. Nat. Commun. 11, 5723 (2020).

